## Supplementary material for "OpenCell: A Low-cost, Open-Source, 3-in-1 device for DNA Extraction": S1 File

**Supporting Information: S1 Figure - OpenCell: A Low-cost, Open-Source, 3-in-1 device for DNA Extraction**

### Contents

**S1 Figure: Wiring diagram** **2**

**S2 Figure: PID Accuracy** **3**

### S1 Figure

##### Wiring Diagram of all electronic components used in OpenCell

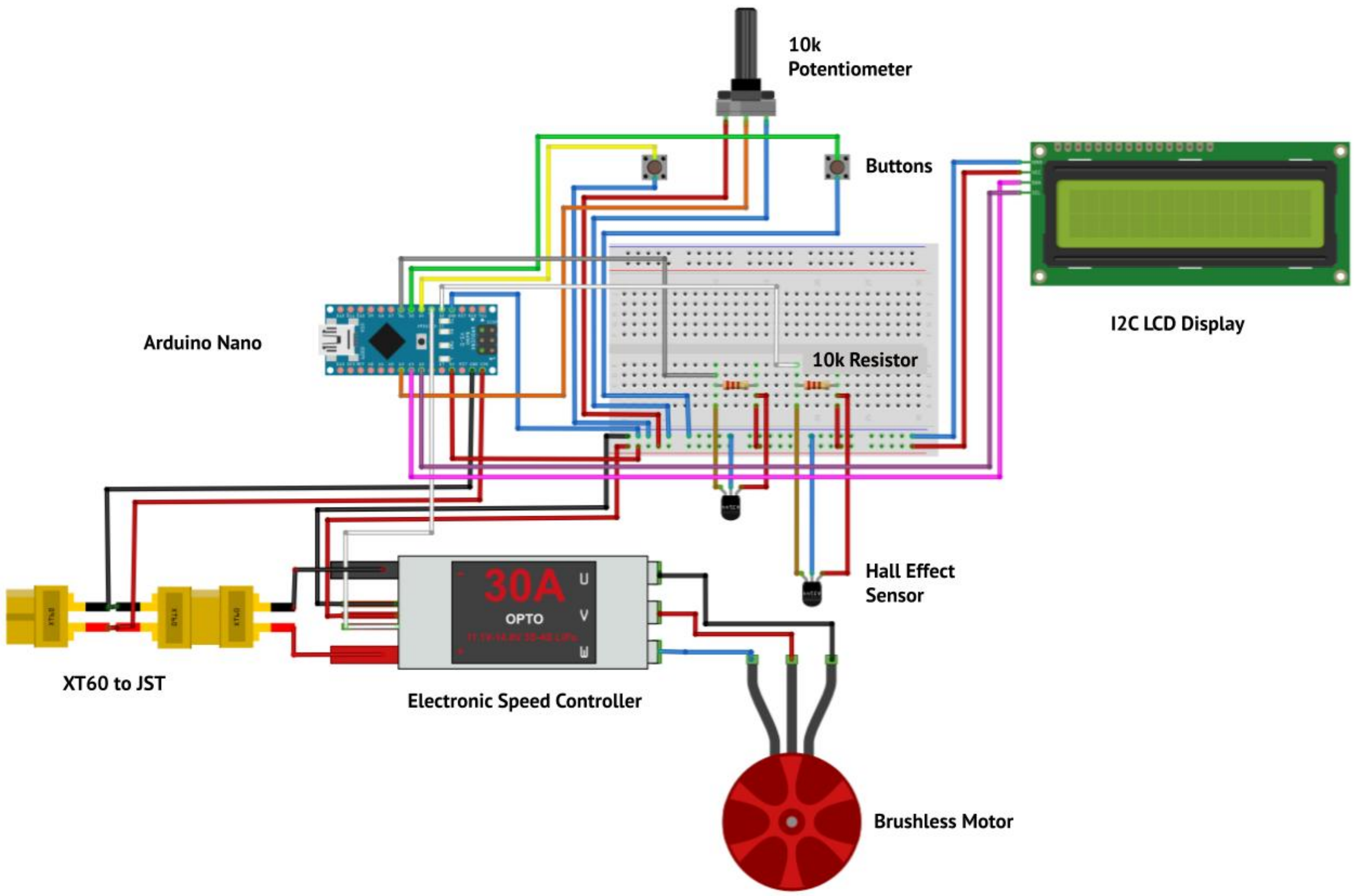

#### S2 Figure

**Accuracy of PID controller over time**

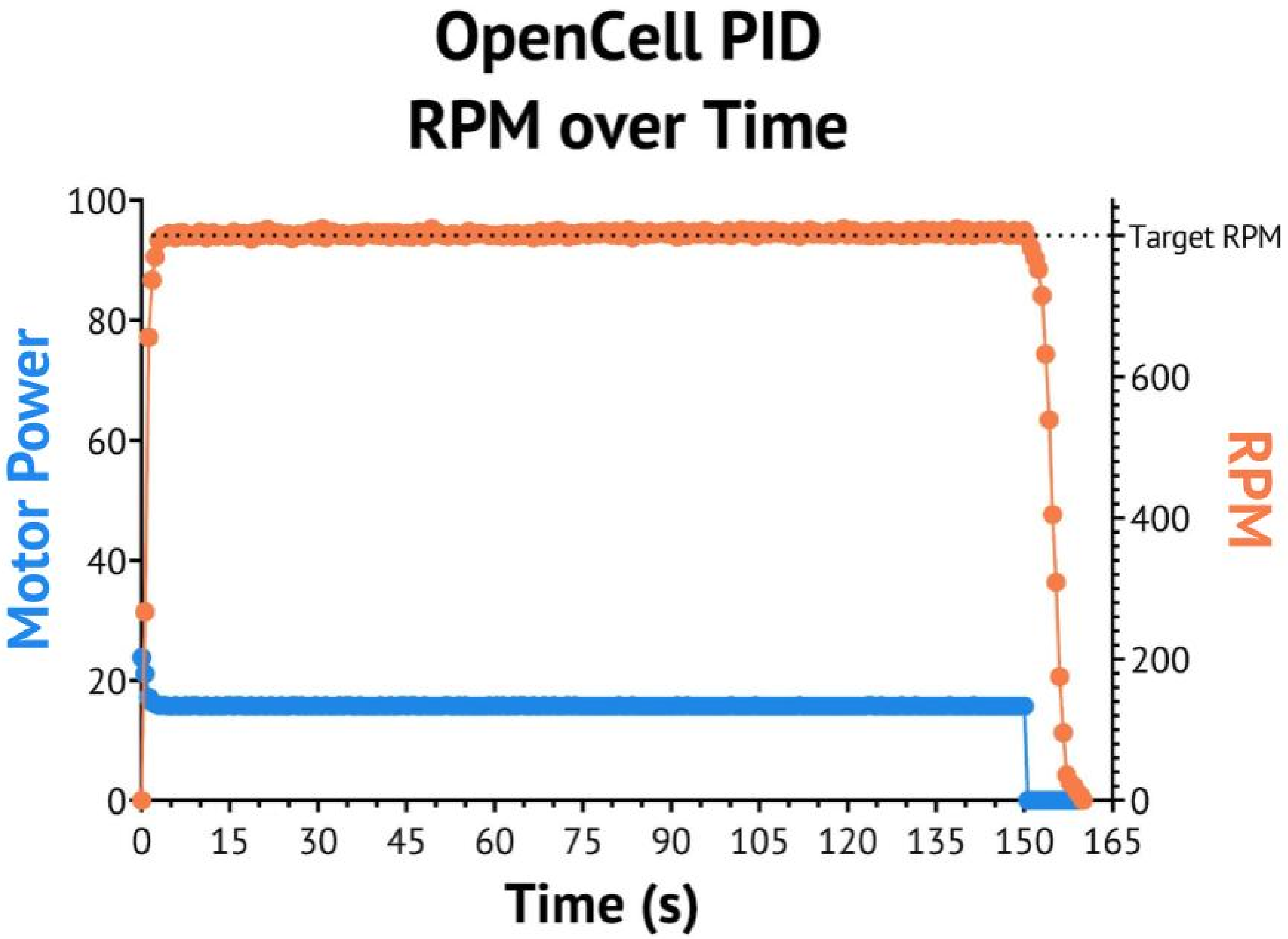
