## Supplementary material for "OpenCell: A Low-cost, Open-Source, 3-in-1 device for DNA Extraction": S1 Table

### Supporting Information: S1 Table - OpenCell: A Low-cost, Open-Source, 3-in-1 device for DNA Extraction

### S1 Table

##### Component Files and Slicer Settings

#### STL Component Files

The following github link contains all necessary STL files for OpenCell: Github Repo

#### 3D Printing Defects

Defects and print defects are a common occurrence in 3D printed components and can substantially impact the strength and performance of printed components. Here are some useful external resource for identifying and correcting for common defects: Simplify3d.com, all3dp.com

#### Slicer Settings

| Component | Layer Height | Perimeters/ Walls | Support | Infill | Special |
|---|---|---|---|---|---|
| **Face** | 0.24 | 3 | **no** | 20% | |
| **Lid** | 0.24 | 5 | **no** | 30% | |
| **Chamber** | 0.24 | 5 | **yes-everywhere** | 30% | |
| **Electronics Lid** | 0.24 | 3 | **no** | 20% | |
| **Upper Arm** | 0.2 | 3 | **yes - touching buildplate** | 20% | |
| **Lower Arm** | 0.2 | 3 | **no** | 20% | |
| **Tertiary Gear** | 0.2 | 5 | **no** | 100% | **Rectilinear Infill** |
| **Secondary Gear** | 0.2 | 5 | **no** | 20% | |
| **Motor Mount** | 0.2 | 3 | **no** | 30% | |
| **Tube Holder** | 0.16 | 3 | **no** | 20% | |
| **Tube Lid** | 0.16 | 3 | **no** | 20% | |
| **Centrifuge** | 0.24 | 3 | **yes - touching buildplate** | 20% | |
| **Button Holders** | 0.24 | 3 | **no** | 20% | |

### Detailed Component Cost Table for OpenCell

| Component | Description | Price | Retail Link |
|---|---|---|---|
| Arduino Nano | Compact microcontroller | $4.25 | [Link] |
| Brushless Motor | 2204 to 2207 2300kv+ Drone Motor | $8.00 | [Link] |
| ESC | Electronic Speed Controller | $10.00 | [Link] |
| Display | i2C LCD Display | $2.69 | [Link] |
| Buttons | 2x Inputs to control OpenCell | $0.21 | [Link] |
| Potentiometer | Input Device to control OpenCell | $0.50 | [Link] |
| Hall Effect Sensor | 2x Magnetic Field Detector | $0.70 | [Link] |
| Small Bearings | 4x 693ZZ bearings | $1.90 | [Link] |
| Large Bearings | 2x 608ZZ bearings | $0.85 | [Link] |
| Breadboard | Base for the circuits | $1.16 | [Link] |
| Wires | Used to make electronic connections | $0.75 | [Link] |
| Magnets | 14 6x3mm magnets | $0.76 | [Link] |
| Fasteners | 4, 8, 10, 12, and 16mm M3 bolts and nuts | $0.35 | [Link] |
| Master Switch | Inline xt60 switch | $2.99 | [Link] |
| DC Power Supply | 6-12v, 2amp DC Power Supply | $9.99 | [Link] |
| XT60 to JST Adapter | Power pass-through for Arduino | $1.97 | [Link] |
| LiPo Battery (Optional) | 3s 11.1V 2200mAh For battery operation | $11.50 | [Link] |
| Filament Cost | Total 425g PLA Filament Cost | $4.25 | |
| **Total** | Excluding batteries | **$49.35** | |

This table provides a detailed breakdown of the cost of each component used in the OpenCell platform. The prices reflect the market costs in 2022. The total cost of the OpenCell platform, excluding the optional LiPo battery, is approximately $47. With the optional LiPo battery and excluding the DC power supply, the total cost is less than $50.
