## Supplementary material for "OpenCell: A Low-cost, Open-Source, 3-in-1 device for DNA Extraction": S2 File: OpenCell_No_PID.ino

```
//OPENCELL NO PID CODE
//BhamlaLab - Georgia Institute of Technology
//Aryan Gupta
//USING CODE ADAPTED FROM https://www.youtube.com/watch?v=u2uJMJWsfsg
/*
  This verion of the code is reccomended for beginner programmers. The run speed must be set manually during operation.
  For those with experience Tuning PID's, see the PID Verison of this Code
*/

#include <Servo.h>
#include <Wire.h>
#include <LiquidCrystal_I2C.h>
#define rightBtn 5
#define leftBtn 4
#define dial 6
#define tach 2
#define motor 3

LiquidCrystal_I2C lcd(0x27, 16, 2);
Servo esc;
int potValue;
unsigned long lastTime_print = 0;


const byte PulsesPerRevolution = 2;  // Set how many pulses there are on each revolution. Default: 2.
const unsigned long ZeroTimeout = 200000;
const byte numReadings = 2;

volatile unsigned long LastTimeWeMeasured;
volatile unsigned long PeriodBetweenPulses = ZeroTimeout + 1000; 
volatile unsigned long PeriodAverage = ZeroTimeout + 1000;
unsigned long FrequencyRaw;
unsigned long FrequencyReal;
unsigned long RPM;
unsigned int PulseCounter = 1;

unsigned long PeriodSum;
unsigned long LastTimeCycleMeasure = LastTimeWeMeasured;

unsigned long CurrentMicros = micros();

unsigned int AmountOfReadings = 1;

unsigned int ZeroDebouncingExtra;


unsigned long readings[numReadings];
unsigned long readIndex;
unsigned long total;
unsigned long average;

void setup()  // Start of setup:
{
  pinMode(rightBtn, INPUT_PULLUP);
  pinMode(leftBtn, INPUT_PULLUP);
  pinMode(dial, INPUT);
  Serial.begin(9600);
  attachInterrupt(digitalPinToInterrupt(tach), Pulse_Event, FALLING);

  lcd.begin();
  lcd.backlight();
  lcd.clear();
  esc.attach(motor, 1000, 2000); 
  esc.write(30);
  delay(1000); 

}  // End of setup.


void loop()  // Start of loop:
{
  homogenize();
}


void updateLCD(int power, int timeLeft) {
  if ((millis() - lastTime_print) >= 500) {
    int pwr = map(power, 1000, 2000, 0, 100);
    lastTime_print = millis();
    lcd.clear();
    lcd.setCursor(0, 0);
    lcd.print("Time: ");
    lcd.setCursor(10, 0);
    lcd.print(timeLeft);
    lcd.setCursor(0, 1);
    lcd.print("RPM:");
    lcd.print(average);
    lcd.setCursor(9, 1);
    lcd.print("P:");
    lcd.print(pwr);
    Serial.print("Period: ");
    Serial.print(PeriodBetweenPulses);
    Serial.print("\tReadings: ");
    Serial.print(AmountOfReadings);
    Serial.print("\tFrequency: ");
    Serial.print(FrequencyReal);
    Serial.print("\tRPM: ");
    Serial.print(RPM);
    Serial.print("\tTachometer: ");
    Serial.println(average);

  }
}

int initializeOpenCell() {
  lcd.clear();
  lcd.setCursor(4, 0);
  lcd.print("OpenCell");
  lcd.setCursor(0, 1);
  lcd.print("Science for All!");
  delay(2000);
}

int startScreen() {
  lcd.clear();
  lcd.setCursor(4, 0);
  lcd.print("OpenCell");
  lcd.setCursor(1, 1);
  lcd.print("Press To Start");
}

int setTime() {
  lcd.clear();
  lcd.setCursor(4, 0);
  lcd.print("Set Time");
  lcd.setCursor(1, 1);
}

int confirmTime(int checkTime) {
  lcd.clear();
  lcd.setCursor(2, 0);
  lcd.print("Confrim Time");
  lcd.setCursor(1, 1);
  lcd.print(checkTime);
}


int runHomogenizer(float duration) {
  lcd.clear();
  lcd.setCursor(0,0);
  lcd.print("Set Dial To Zero");
  delay(3000);
  float msTime = duration * 1000 * 60;
  long startTime = millis();
  long endTime = startTime + msTime;
  int timeElapsed = 0;
  int printTime = 0;
  while (millis() < endTime) {
    if(digitalRead(5) == 0) endTime = 0; 
    potValue = analogRead(A3);
    potValue = map(potValue, 0, 1023, 1000, 2000);
    if(digitalRead(6) < 1) esc.write(potValue );
    else esc.writeMicroseconds(0);
    printTime = (endTime - millis()) / 1000;
    updateLCD(potValue, printTime);
    checkSpeed();
  }
  esc.write(0);
  lcd.clear();
  lcd.setCursor(2, 0);
  lcd.print("OpenCell Pro");
  lcd.setCursor(1, 1);
  lcd.print("Finished");
  delay(1000);
}


int homogenize() {
  int  pwmOut = 0;
  bool timeSet = false;
  bool timeConfirm = false;
  float runTime;
  lcd.begin();
  lcd.backlight();
  lcd.setCursor(1, 0);
  initializeOpenCell();
  startScreen();
  //SET RUN TIME
  while (digitalRead(rightBtn) == 1) {}
  setTime();
  delay(1000);
  while (timeSet == false) {
    potValue = analogRead(A3);
    potValue = map(potValue,0,1023, 0, 80);
    float timer = potValue / 4.00;
    lcd.setCursor(5, 1);
    lcd.print(timer);
    if (digitalRead(rightBtn) == 0) {
      runTime = timer;
      timeSet = true;
    }
  }
  delay(1000);
  //  confirmTime(runTime);
  //  while(rightButton ==1){
  //    rightButton = digitalRead(4);
  //    leftButton = digitalRead(5);
  //  }

  //RUN HOMOGENIZER
  runHomogenizer(runTime);
}

void checkSpeed() {
  LastTimeCycleMeasure = LastTimeWeMeasured;
  CurrentMicros = micros();
  if (CurrentMicros < LastTimeCycleMeasure)
  {
    LastTimeCycleMeasure = CurrentMicros;
  }

  FrequencyRaw = 10000000000 / PeriodAverage;

  if (PeriodBetweenPulses > ZeroTimeout - ZeroDebouncingExtra || CurrentMicros - LastTimeCycleMeasure > ZeroTimeout - ZeroDebouncingExtra)
  {
    FrequencyRaw = 0;
    ZeroDebouncingExtra = 2000;
  }
  else
  {
    ZeroDebouncingExtra = 0;
  }

  FrequencyReal = FrequencyRaw / 10000;
  RPM = FrequencyRaw / PulsesPerRevolution * 60;
  RPM = RPM / 10000; 


  total = total - readings[readIndex];
  readings[readIndex] = RPM;
  total = total + readings[readIndex];
  readIndex = readIndex + 1;

  if (readIndex >= numReadings) readIndex = 0;

  // Calculate the average:
  average = total / numReadings;

}


void Pulse_Event()  // The interrupt runs this to calculate the period between pulses:
{
  PeriodBetweenPulses = micros() - LastTimeWeMeasured; 
  LastTimeWeMeasured = micros();

  if (PulseCounter >= AmountOfReadings)
  {
    PeriodAverage = PeriodSum / AmountOfReadings;
    PulseCounter = 1;
    PeriodSum = PeriodBetweenPulses;

    int RemapedAmountOfReadings = map(PeriodBetweenPulses, 40000, 5000, 1, 10);
    RemapedAmountOfReadings = constrain(RemapedAmountOfReadings, 1, 10);
    AmountOfReadings = RemapedAmountOfReadings;
  }
  else
  {
    PulseCounter++;
    PeriodSum = PeriodSum + PeriodBetweenPulses;
  }

}
```
