## Supplementary material for "OpenCell: A Low-cost, Open-Source, 3-in-1 device for DNA Extraction": S2 File: OpenCell_PID.ino

```
//OPENCELL PID CODE
//BhamlaLab - Georgia Institute of Technology
//Aryan Gupta

//USING CODE ADAPTED FROM https://www.youtube.com/watch?v=u2uJMJWsfsg
/*
  This verion of the code is reccomended for experienced programmers. A PID allows for accurate and precise control.
  A PID must be tuned and improper implementation can lead to instability, component damage, and overheating.
  We reccomend a low Kp value and a slightly overdamped system.
*/

#include <Servo.h>
#include <Wire.h>
#include <LiquidCrystal_I2C.h>
#define rightBtn 5
#define leftBtn 4
#define dial A3
#define tach 2
#define motor 3
#define lidSensor 6
float Kp = 0;
float Ki = 0;
float Kd = 0;
PID 
LiquidCrystal_I2C lcd(0x27, 16, 2);
Servo esc;
int potValue;
unsigned long lastTime_print = 0;


const byte PulsesPerRevolution = 2;  // Set how many pulses there are on each revolution. Default: 2.
const unsigned long ZeroTimeout = 200000;
const byte numReadings = 2;

volatile unsigned long LastTimeWeMeasured;
volatile unsigned long PeriodBetweenPulses = ZeroTimeout + 1000; 
volatile unsigned long PeriodAverage = ZeroTimeout + 1000;
unsigned long FrequencyRaw;
unsigned long FrequencyReal;
unsigned long RPM;
unsigned int PulseCounter = 1;

unsigned long PeriodSum;
unsigned long LastTimeCycleMeasure = LastTimeWeMeasured;

unsigned long CurrentMicros = micros();

unsigned int AmountOfReadings = 1;

unsigned int ZeroDebouncingExtra;


unsigned long readings[numReadings];
unsigned long readIndex;
unsigned long total;
unsigned long average;

void setup()  // Start of setup:
{
  pinMode(rightBtn, INPUT_PULLUP);
  pinMode(leftBtn, INPUT_PULLUP);
  pinMode(dial, INPUT);
  Serial.begin(9600);
  attachInterrupt(digitalPinToInterrupt(tach), Pulse_Event, FALLING);

  lcd.begin();
  lcd.backlight();
  lcd.clear();
  esc.attach(motor, 1000, 2000); 
  esc.write(30);
  delay(1000); 

}  // End of setup.


void loop()  // Start of loop:
{
  homogenize();
}


void updateLCD(int power, int timeLeft) {
  if ((millis() - lastTime_print) >= 500) {
    int pwr = map(power, 1000, 2000, 0, 100);
    lastTime_print = millis();
    lcd.clear();
    lcd.setCursor(0, 0);
    lcd.print("Time: ");
    lcd.setCursor(10, 0);
    lcd.print(timeLeft);
    lcd.setCursor(0, 1);
    lcd.print("RPM:");
    lcd.print(average);
    lcd.setCursor(9, 1);
    lcd.print("P:");
    lcd.print(pwr);
    Serial.print("Period: ");
    Serial.print(PeriodBetweenPulses);
    Serial.print("\tReadings: ");
    Serial.print(AmountOfReadings);
    Serial.print("\tFrequency: ");
    Serial.print(FrequencyReal);
    Serial.print("\tRPM: ");
    Serial.print(RPM);
    Serial.print("\tTachometer: ");
    Serial.println(average);

  }
}

int initializeOpenCell() {
  lcd.clear();
  lcd.setCursor(4, 0);
  lcd.print("OpenCell");
  lcd.setCursor(0, 1);
  lcd.print("Science for All!");
  delay(2000);
}

int startScreen() {
  lcd.clear();
  lcd.setCursor(4, 0);
  lcd.print("OpenCell");
  lcd.setCursor(1, 1);
  lcd.print("Press To Start");
}

int setTime() {
  lcd.clear();
  lcd.setCursor(4, 0);
  lcd.print("Set Time");
  lcd.setCursor(1, 1);
}

int confirmTime(int checkTime) {
  lcd.clear();
  lcd.setCursor(2, 0);
  lcd.print("Confirm Time");
  lcd.setCursor(1, 1);
  lcd.print(checkTime);
}

int setSpeed() {
  lcd.clear();
  lcd.setCursor(4, 0);
  lcd.print("Set Speed");
  lcd.setCursor(1, 1);
}

int confirmSpeed(int checkSpeed) {
  lcd.clear();
  lcd.setCursor(2, 0);
  lcd.print("Confirm Speed");
  lcd.setCursor(1, 1);
  lcd.print(checkSpeed);
}


int runHomogenizer(float duration, float runSpeed) {
  lcd.clear();
  lcd.setCursor(0,0);
  lcd.print("Set Dial To Zero");
  delay(3000);
  float msTime = duration * 1000 * 60;
  long startTime = millis();
  long endTime = startTime + msTime;
  int timeElapsed = 0;
  int printTime = 0;
  float error = 0;
  float cumulativeError = 0;
  float deltaError = 0;
  float lastError = 0;
  float outputValue = 0;
  while (millis() < endTime) {
    error = runSpeed - average;
    cumulativeError += error;
    deltaError = error - lastError;
    lastError = error;
    outputValue = Kp*error + Ki*cumulativeError + Kd*deltaError;
    if(outputValue < 1000) outputValue = 1000;
    if(outputValue > 2000) outputValue = 2000;
    if((digitalRead(rightBtn) == 0) || (digitalRead(leftBtn) == 0)){
      esc.write(0);
      endTime = 0; 
    }
    if(digitalRead(lidSensor) < 1) {
      esc.write(outputValue);
    }
    else{
       esc.writeMicroseconds(0);
       endTime = 0;
    }

    printTime = (endTime - millis()) / 1000;
    updateLCD(runSpeed, printTime);
    checkSpeed();
  }
  esc.write(0);
  lcd.clear();
  lcd.setCursor(2, 0);
  lcd.print("OpenCell Pro");
  lcd.setCursor(1, 1);
  lcd.print("Finished");
  delay(1000);
}


int homogenize() {
  int  pwmOut = 0;
  bool timeSet = false;
  bool speedSet = false;
  float runTime;
  float runSpeed;
  
  lcd.begin();
  lcd.backlight();
  lcd.setCursor(1, 0);
  initializeOpenCell();
  startScreen();
  //SET RUN TIME
  while (digitalRead(rightBtn) == 1) {}
  setTime();
  delay(1000);
  while (timeSet == false) {
    potValue = analogRead(A3);
    potValue = map(potValue,0,1023, 0, 80);
    float timer = potValue / 4.00;
    lcd.setCursor(5, 1);
    lcd.print(timer);
    if (digitalRead(rightBtn) == 0) {
      runTime = timer;
      timeSet = true;
    }
  }
  delay(1000);
  confirmTime(runTime);
  while(digitalRead(rightBtn) ==1){}

  //SET RUN SPEED
  delay(1000);
  setSpeed();
  delay(1000);
  while (speedSet == false) {
    potValue = analogRead(A3);
    potValue = map(potValue,0,1023, 200, 5000);
    float speed = potValue / 4.00;
    lcd.setCursor(5, 1);
    lcd.print(speed);
    if (digitalRead(rightBtn) == 0) {
      runSpeed = speed;
      speedSet = true;
    }
  }
  delay(1000);
  confirmTime(runTime);
  while(digitalRead(rightBtn) ==1){}

  //RUN HOMOGENIZER
  runHomogenizer(runTime, runSpeed);
}

void checkSpeed() {
  LastTimeCycleMeasure = LastTimeWeMeasured;
  CurrentMicros = micros();
  if (CurrentMicros < LastTimeCycleMeasure)
  {
    LastTimeCycleMeasure = CurrentMicros;
  }

  FrequencyRaw = 10000000000 / PeriodAverage;

  if (PeriodBetweenPulses > ZeroTimeout - ZeroDebouncingExtra || CurrentMicros - LastTimeCycleMeasure > ZeroTimeout - ZeroDebouncingExtra)
  {
    FrequencyRaw = 0;
    ZeroDebouncingExtra = 2000;
  }
  else
  {
    ZeroDebouncingExtra = 0;
  }

  FrequencyReal = FrequencyRaw / 10000;
  RPM = FrequencyRaw / PulsesPerRevolution * 60;
  RPM = RPM / 10000; 


  total = total - readings[readIndex];
  readings[readIndex] = RPM;
  total = total + readings[readIndex];
  readIndex = readIndex + 1;

  if (readIndex >= numReadings) readIndex = 0;

  // Calculate the average:
  average = total / numReadings;

}


void Pulse_Event()  // The interrupt runs this to calculate the period between pulses:
{
  PeriodBetweenPulses = micros() - LastTimeWeMeasured; 
  LastTimeWeMeasured = micros();

  if (PulseCounter >= AmountOfReadings)
  {
    PeriodAverage = PeriodSum / AmountOfReadings;
    PulseCounter = 1;
    PeriodSum = PeriodBetweenPulses;

    int RemapedAmountOfReadings = map(PeriodBetweenPulses, 40000, 5000, 1, 10);
    RemapedAmountOfReadings = constrain(RemapedAmountOfReadings, 1, 10);
    AmountOfReadings = RemapedAmountOfReadings;
  }
  else
  {
    PulseCounter++;
    PeriodSum = PeriodSum + PeriodBetweenPulses;
  }

}
```
