## Supplementary material for "OpenCell: A Low-cost, Open-Source, 3-in-1 device for DNA Extraction": S3 File

**Supporting Information: S1 Figure - OpenCell: A Low-cost, Open-Source, 3-in-1 device for DNA Extraction**

### Contents

**Open Cell Operating Instructions** **2**

**OpenCell DNA Extraction Procedure** **3**

**OpenCell Optimization** **4**

**DNA extraction protocol** **6**

**OpenCell Assembly Instructions** **7**

### Operating Instructions

#### OpenCell General Operating Instructions

##### Initial Instructions

Power on the device by plugging in the 6v power supply on the left side port and connecting the blue cable to the right side port of the device. The LCD Display should turn on and text should appear on the screen. At the same time, you should hear short beeps to signify the Motor is working properly. Place the attachment you are using securely to the mounting point using the magnets, making sure that the attachment is level.

##### Bead Homogenization

After completing the initial instructions, place the homogenization attachment onto the mounting point, making sure the attachment is level. During this step, be sure to line up the tube holders so that they are in the same orientation. Insert the sample tubes into the tube holders and make sure they are in the same orientation with the lids pointing in the same direction. Firmly press them into place. Then slide the cover into place. The LCD display will prompt you to ”Press to Start.” press the left button to proceed. When prompted to set the speed, use the dial to set your desired speed and press the left button to confirm. The LCD will then display the run time, RPM, and ”Power.” The homogenization run has begun. To start the cycle turn the dial to change the amount of power inflow (and thus RPM).

##### Centrifugation

After completing the initial instructions, place the centrifuge attachment onto the mounting point, making sure the attachment is level. Insert the sample tubes into the tube holders. Firmly press them into place making sure to balance the tubes. For example if processing 2 tubes place them opposite each other, if processing 3 tubes, alternate to create equal spaces. If processing only 1 tube, use a balancing tube filled with approximately the same amount of liquid to offset the single sample. The LCD display will prompt you to ”Press to Start.” press the left button to proceed. When prompted to set the speed, use the dial to set your desired speed and press the left button to confirm. The LCD will then display the run time, RPM, and ”Power.” The homogenization run has begun. Turn the dial to change the amount of power inflow (and thus RPM).

### OpenCell DNA Extraction Procedure

**Personal Protective Equipment Notice:** Please wear a lab coat, safety goggles, and some form of hearing protection whenever OpenCell is being operated. Nitrile gloves should be worn with this protocol, and other appropriate PPE should be worn for any downstream applications.

1) Add 5–100 mg of fresh or frozen plant tissue and 500 µl of Solution CD1 to a 2 ml tissue disruption tube. Mix the sample using the OpenCell Lysis module at 500 RPM for 5 seconds. Note: If your sample is high in phenolic compounds, add 450 µl Solution CD1 and 50 µl Solution PS.
2) Homogenize for 3 minutes at 850 RPM using the OpenCell Cell Lysis Attachment.
3) Centrifuge the tissue disruption tubes at 6000 RPM (3000 RCF) for 5 min.
4) Transfer the supernatant to a clean 1.5 ml microcentrifuge tube (provided).
   - Note: Expect 350–450 µl. The supernatant may still contain some plant particles.
5) Add 200 µl Solution CD2 and mix using Cell Lysis Attachment at 500 RPM for 5 seconds Note:
   For problematic samples, add 250 µl Solution CD2.
6) Centrifuge at 6000 RPM (3000 RCF) for 3 min at room temperature. Avoiding the pellet, transfer the supernatant to a clean 1.5 ml microcentrifuge tube (provided). - Note: Expect 400–500 µl.
7) Add 500 µl of Buffer APP and mix using Cell Lysis Attachment at 500 RPM for 5 seconds.
8) Load 600 µl lysate onto an MB Spin Column. Centrifuge at 6000 RPM (3000 RCF) for 3 min.
9) Discard the flow-through and repeat step 8 to ensure that all of the lysate has passed through the MB spin column.
10) Place the MB spin column into a clean 2 ml collection tube.
11) Add 650 µl Buffer AW1 to the MB spin column. Centrifuge at 6000 RPM (3000 RCF) for 3 min. Discard the flow-through and place the MB spin column back into the same 2 ml collection tube.
12) Add 650 µl of Buffer AW2 to the MB spin column. Centrifuge at 6000 RPM (3000 RCF) for 3 min. Discard the flow-through and place the MB spin column into the same 2 ml collection tube. 13) Centrifuge at up to 6000 RPM (3000 RCF) for 5 min. Place the MB spin column into a new 1.5 ml elution tube (provided).
14) Add 50–100 µl of Buffer EB to the center of the white filter membrane.
15) Centrifuge at 6000 RPM (3000 RCF) for 3 min. Discard the MB spin column. The DNA is now ready for downstream application

### Detailed Optimization and Characterization of OpenCell

#### Detailed Procedure for DNA Extraction

The DNA extraction process involves four main steps: lysis, binding, washing, and elution. During the lysis step, a bead-mill (cell lysis module) is used in conjunction with chemical detergents to rupture cell membranes and expose intracellular materials. The binding step first uses a vortex mixer to accelerate a precipitation reaction that removes extraneous cellular debris from the lysate. A centrifuge is then used to pass the resulting solution through a silica-based membrane, which selectively binds to DNA. The wash steps use a centrifuge to push a salt/ethanol solution through the membrane, removing any remaining contaminants from the silica medium. Finally, a centrifuge is required once again to elute the DNA from the membrane using a buffer.

#### Considerations for OpenCell Optimization

Operational speeds and times in published DNA extraction protocols assume the use of commercially available and calibrated equipment. As a result, properly contextualizing the performance of the OpenCell device in relation to these protocols is crucial for reliably achieving high-quality results. When using the bead-mill attachment for cell lysis, it is important to select an operational speed that provides sufficient energy to effectively rupture cell membranes and release intracellular materials. Low speeds may not generate enough force for complete lysis, while high speeds can lead to excessive shearing of DNA. Likewise, an insufficient run duration will lyse an insufficient amount of DNA for proper study, while an excessive run duration will generate significantly more heat and risk damaging DNA and the device itself. These factors must be properly balanced for both the OpenCell bead-mill and centrifuge modules.

#### Cell Lysis Module Characterization

In our characterization of the cell lysis module, we conducted two experiments. The first experiment investigated the impact of operational speed on DNA yield and quality while maintaining a constant run duration. The second experiment examined the effect of operational duration on DNA yield and quality while keeping the operational speed constant.

We established an operating range of speeds for the cell lysis module, spanning from 425 to 1000 RPM. This range was subdivided into five groups - 425 RPM, 550 RPM, 725 RPM, 850 RPM, and 1000 RPM - each processed for 120 seconds with three replicates per group.

Our results indicate that speeds at and below 550 RPM do not generate enough energy per collision to effectively lyse cell walls, thus producing little usable DNA. Past 550 RPM, DNA yields increased dramatically with diminishing returns after 725 and 850 RPM. DNA purity measurements showed no significant improvement beyond 725 RPM. Based on these findings, we determined an optimal operating speed range of 725-850 RPM for the cell lysis module.

##### Optimization of Operational Duration

While optimizing OpenCell's operational run-time, we noticed that sample tubes were warm to the touch (30+deg Celsius) after 5 minutes of operation. Additionally, the cell lysis mechanism exerted a considerable load on the OpenCell power train, causing the ESC and DC Motor to exceed 40 $^{\circ}C$ after 5 minutes of continuous operation. Prolonged operation at this temperature could potentially damage these components

and increase the risk of deformation in the low-temperature PLA thermoplastics used in the construction of OpenCell's chassis.
Considering these factors, the second experimental group was subdivided into five sections - 30 seconds, 60 seconds, 120 seconds, 180 seconds, and 300 seconds. For this particular experiment, the operational speed was fixed to 725 RPM.
The results of the time characterization experiment show a similar trend. Run times below 60 seconds were unable to produce a viable DNA yield. However, DNA yield increased significantly between 60 and 180 seconds, with diminishing returns beyond 180 seconds. For runs longer than 60 seconds, there was no appreciable improvement in DNA purity. From this data, we determined an ideal operating duration range of 120-180 seconds for the bead-mill attachment.

##### Centrifugation Module Characterization

We conducted a simplified experiment to characterize the centrifugation module. Because the system can only produce a maximum of 3000 RCF, well below the protocol's suggested 10,000 RCF-12,000 RCF, there is little value in modulating the speed parameter. The remaining factor that must be optimized is the operation time.
Centrifugation is required in multiple steps within a DNA extraction protocol, with centrifugation duration varying significantly between each step. For this reason, our experimental groups were divided into five groups - 0.5x, 1x, 1.5x, 2x, and 2.5x - where each group represents a time factor applied to the protocol recommended duration for a given step.
The results of the centrifuge characterization experiment indicate that centrifugation time has a lesser effect on DNA yield than on the A260/A280 ratio. Our data shows a gradual increase in both DNA yield and A260/A280 ratio between groups 0.5x to 1.5x, with no appreciable increase in either beyond a factor of 1.5x. Although all groups were able to produce viable DNA yields, factors below 1.5x were unable to reach an A260/280 ratio of 1.8, implying that the DNA was not fully isolated. Thus, we concluded that 1.5x represents the most efficient factor for centrifugation time.

#### Donwstream Validation Procedures

We target the Ribulose bisphosphate carboxylase large chain (Rbcl) gene found natively in *Spinacia oleracea* for downstream analysis. With the DNA extracted from the contextualization experiments in the prior section, we perform a Polymerase Chain Reaction (PCR) using the New England BioLabs Taq 5X Master Mix protocol and primers purchased from Carolina Biological. We purify the PCR product using the Promega Wizard SV Gel and PCR Cleanup System. We analyze the purified product using gel electrophoresis. The gel is run on a 2% agarose gel at 220V for 20 minutes, and the DNA is stained with SYBR Safe.

### The DNA extraction protocol provided with the commercially available Qiagen Plant Pro Kit

1) Add 5–100 mg of fresh or frozen plant tissue and 500 µl of Solution CD1 to a 2 ml tissuedisruption tube. Vortex briefly to mix. - Note: If your sample is high in phenolic compounds, add 450 µl Solution CD1 and 50 µl Solution PS.
2) Homogenize using one of these methods:

   **a) Vortex:** Secure tissue disruption tubes to a Vortex Adapter (cat. no. 13000-V1-24) and vortex at maximum speed for 10 min.

   **b) TissueLyser II:** Most plant samples can be lysed with the TissueLyser II, using the TissueLyser Adapter Set 2 x 24: Place samples in the TissueLyser II and run at 24 Hz for 2 min. Reorient the adapter so the side closest to the machine body becomes furthest from it, and then run the TissueLyser again at 24 Hz for another 2 min.

   **c) PowerLyzer® 24 Homogenizer:** Tissue disruption tubes must be properly balanced in the tube holder of the PowerLyzer 24 Homogenizer. Homogenize the tissue for 1 cycle at the appropriate speed depending on sample type for 2 min.

3) Centrifuge the tissue disruption tubes at 12,000 x g for 2 min.
4) Transfer the supernatant to a clean 1.5 ml microcentrifuge tube (provided). - Note: Expect 350–450 µl. The supernatant may still contain some plant particles.
5) Add 200 µl Solution CD2 and vortex for 5 s. - Note: For problematic samples, add 250 µl Solution CD2.
6) Centrifuge at 12,000 x g for 1 min at room temperature. Avoiding the pellet, transfer the supernatant to a clean 1.5 ml microcentrifuge tube (provided). - Note: Expect 400–500 µl.
7) Add 500 µl of Buffer APP and vortex for 5 s.
8) Load 600 µl lysate onto an MB Spin Column. Centrifuge at 12,000 x g for 1 min.
9) Discard the flow-through and repeat step 8 to ensure that all of the lysate has passed through the MB spin column.
10) Place the MB spin column into a clean 2 ml collection tube (provided).
11) Add 650 µl Buffer AW1 to the MB spin column. Centrifuge at 12,000 x g for 1 min. Discard the flow-through and place the MB spin column back into the same 2 ml collection tube.
12) Add 650 µl of Buffer AW2 to the MB spin column. Centrifuge at 12,000 x g for 1 min. Discard the flow-through and place the MB spin column into the same 2 ml collection tube.
13) Centrifuge at up to 16,000 x g for 2 min. Place the MB spin column into a new 1.5 ml elution tube (provided).
14) Add 50–100 µl of Buffer EB to the center of the white filter membrane.
15) Centrifuge at 12,000 x g for 1 min. Discard the MB spin column. The DNA is now ready for downstream applications.

### Instructions for Assembly
### Lysis Attachment

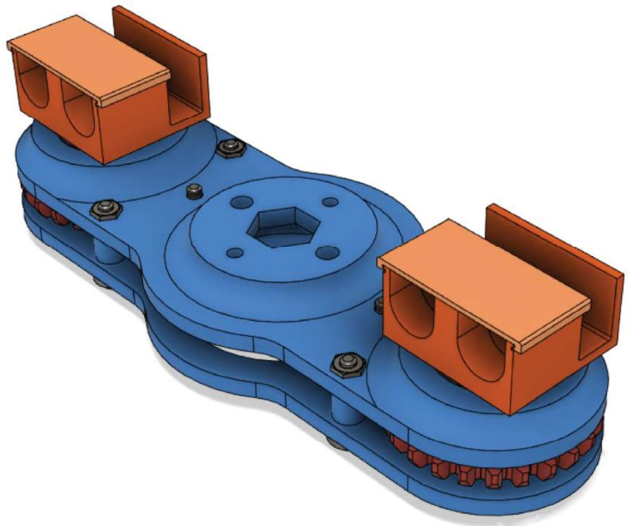

| | | | |
|---|---|---|---|
| 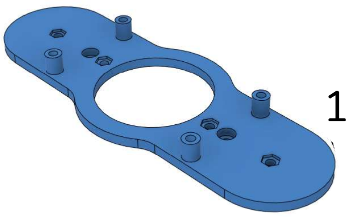 1 x | 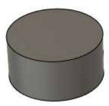 4 x | 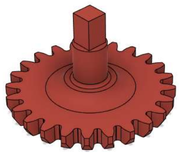 2 x | 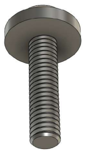 2 x m3x12mm |
| 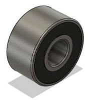 2 x small bearing | 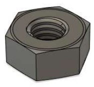 8 x | 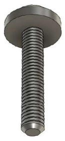 6 x m3x16mm | 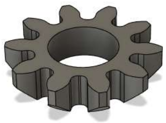 2 x |
| 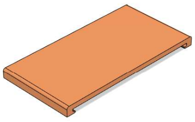 2 x | 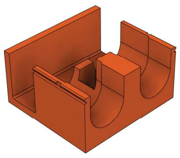 2 x | 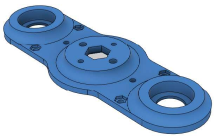 1 x | 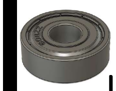 2 x large bearing |

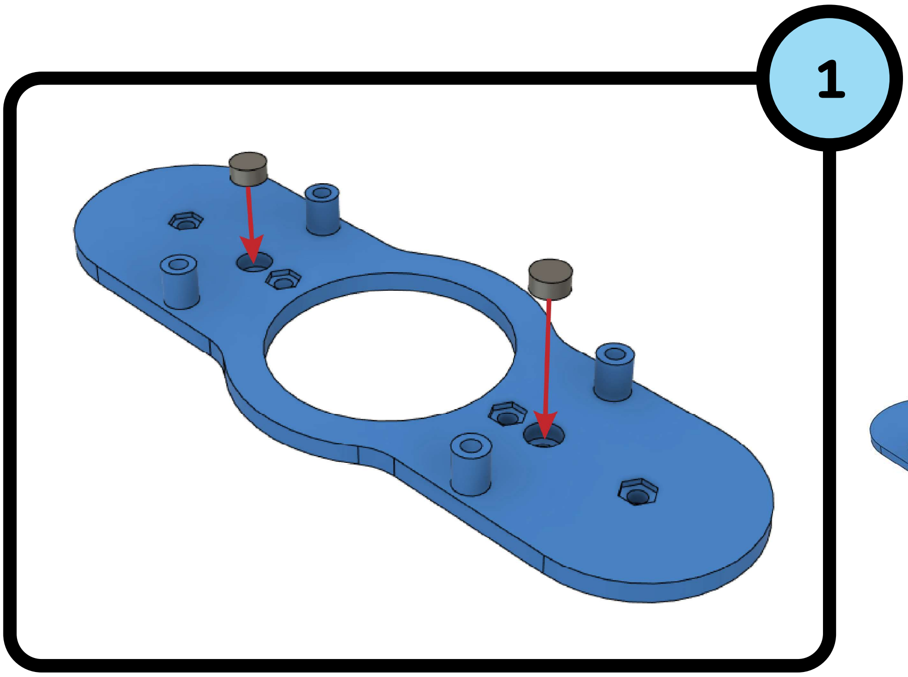


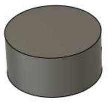

2 x

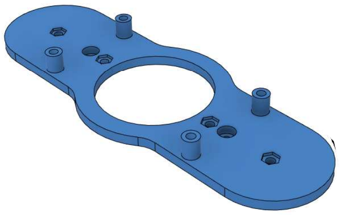

1 x

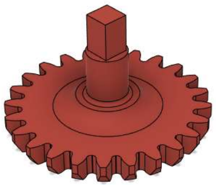

2 x

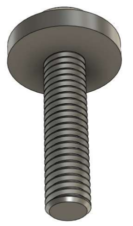

2 x
m3x12mm

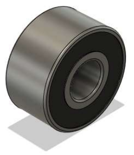

2 x
small bearing

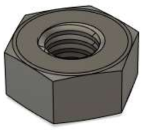

2 x

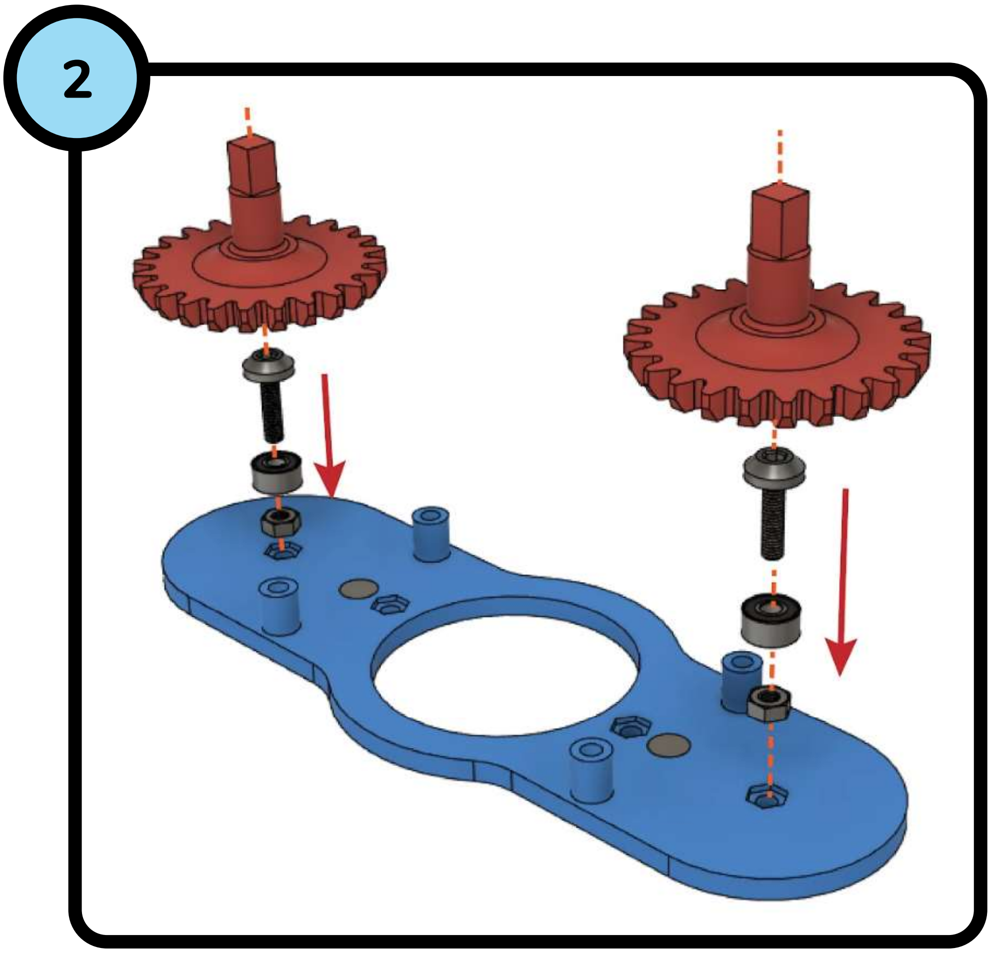

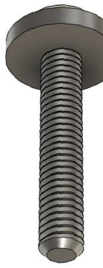

2 x
m3x16mm

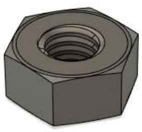

2 x

## 3

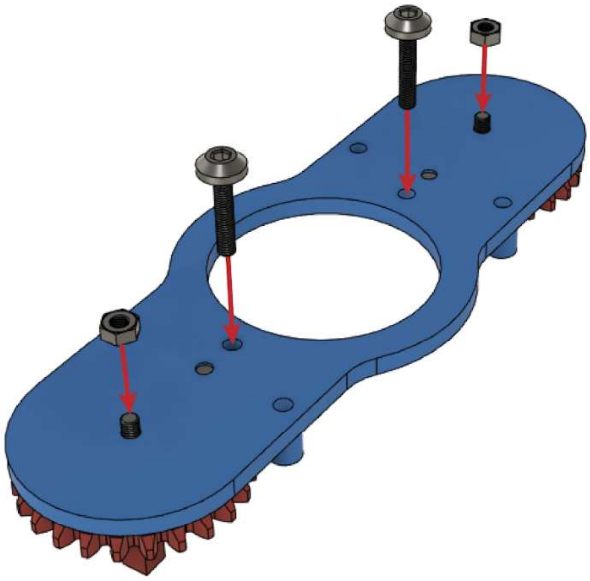

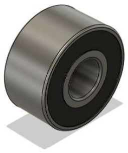

2 x
small bearing

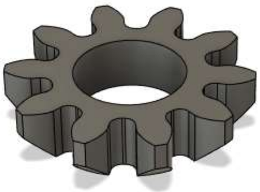

2 x

## 4

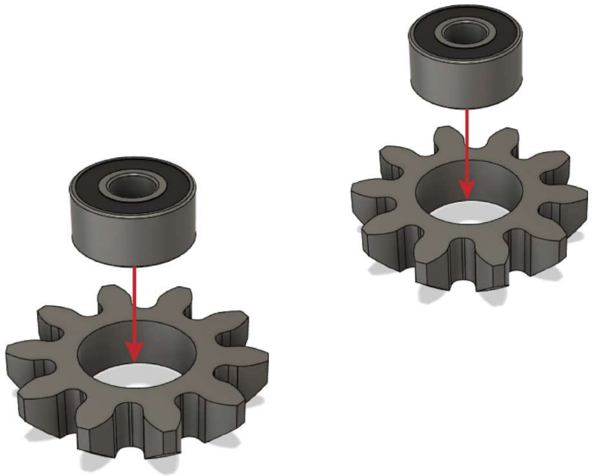

## 5

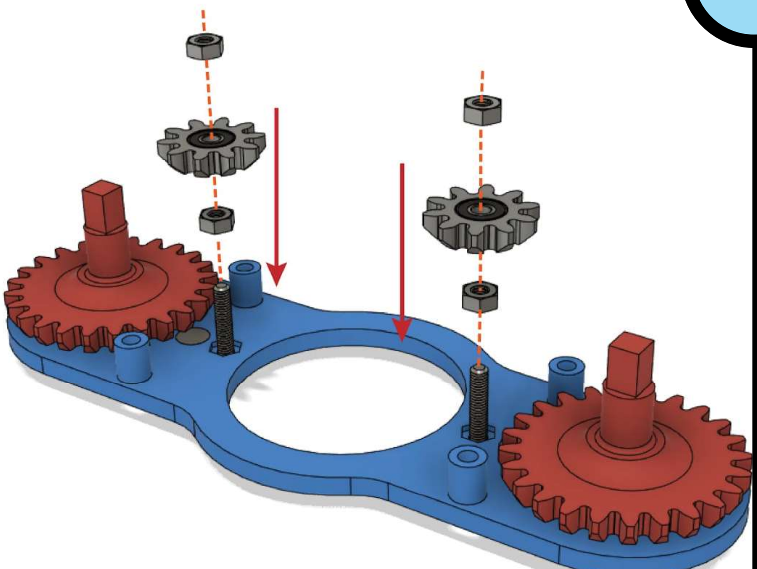

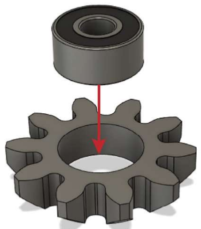

2 x

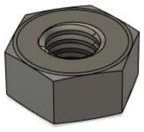

4 x

## 6

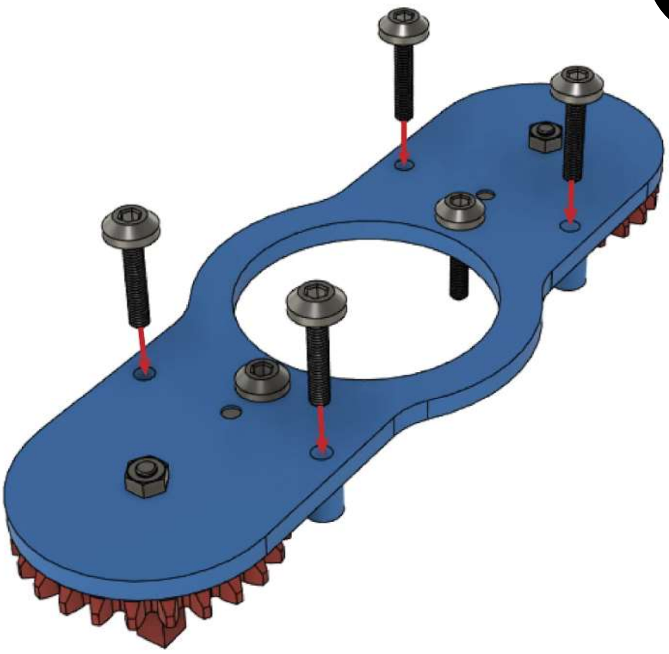

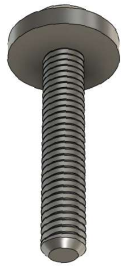

4 x
m3x16mm

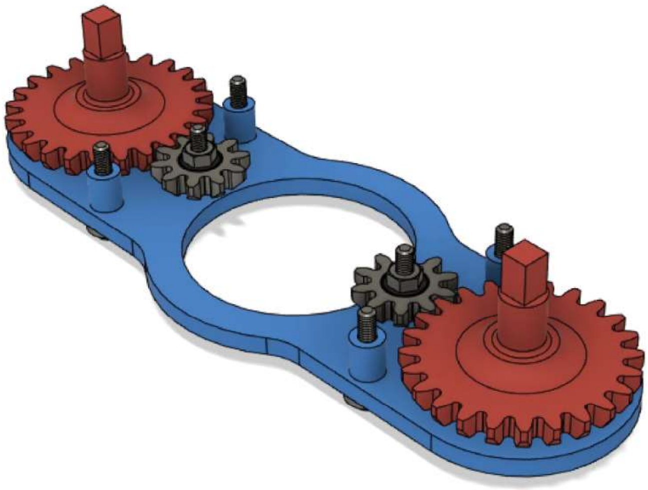

## 7

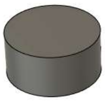

2 x

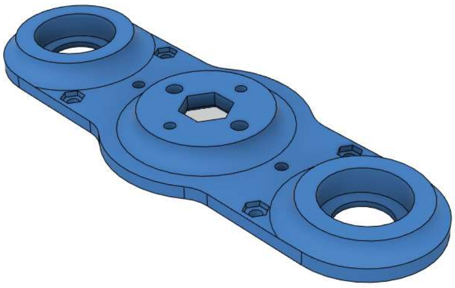

1 x

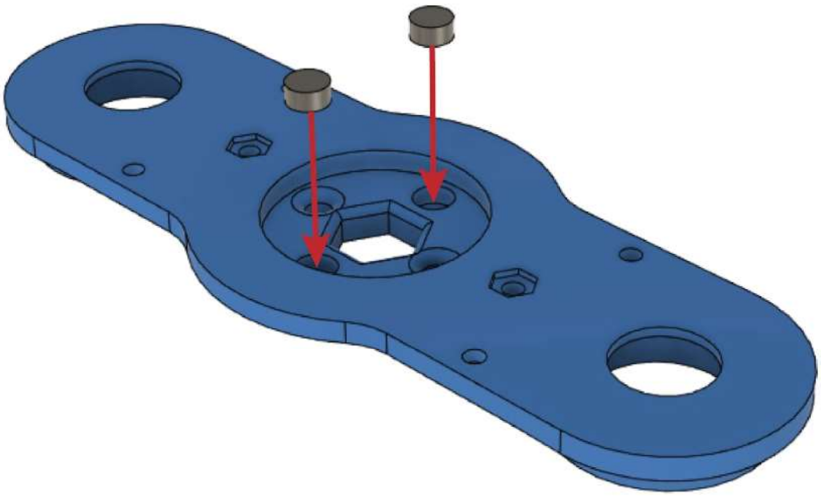

**8**

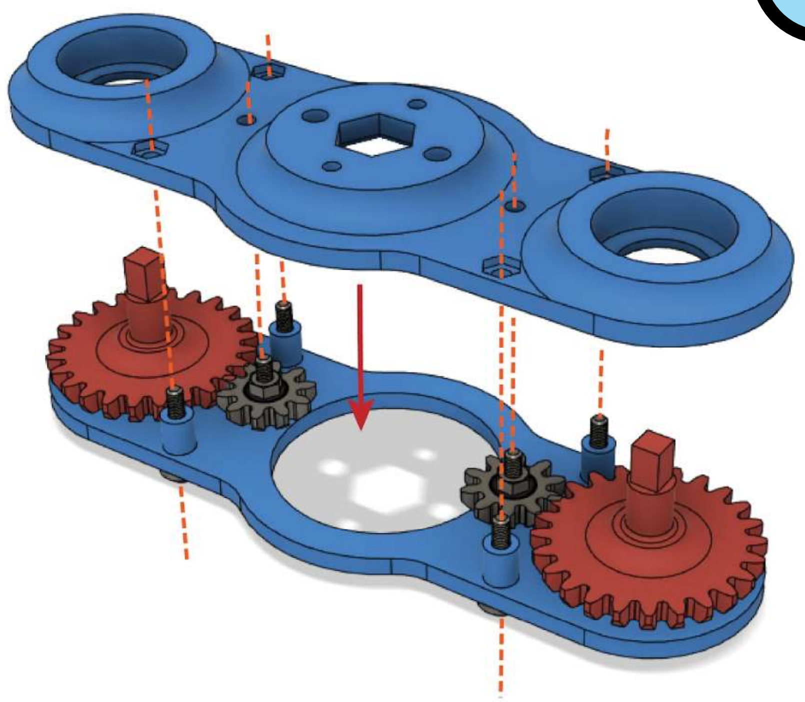

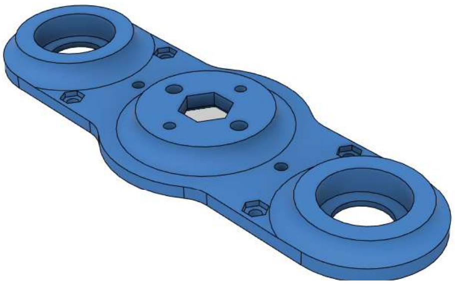

1 x

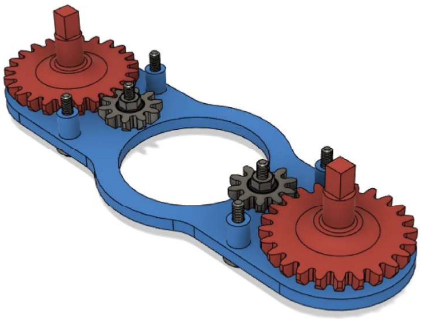

1 x

**9**

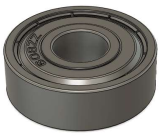

2 x
large bearing

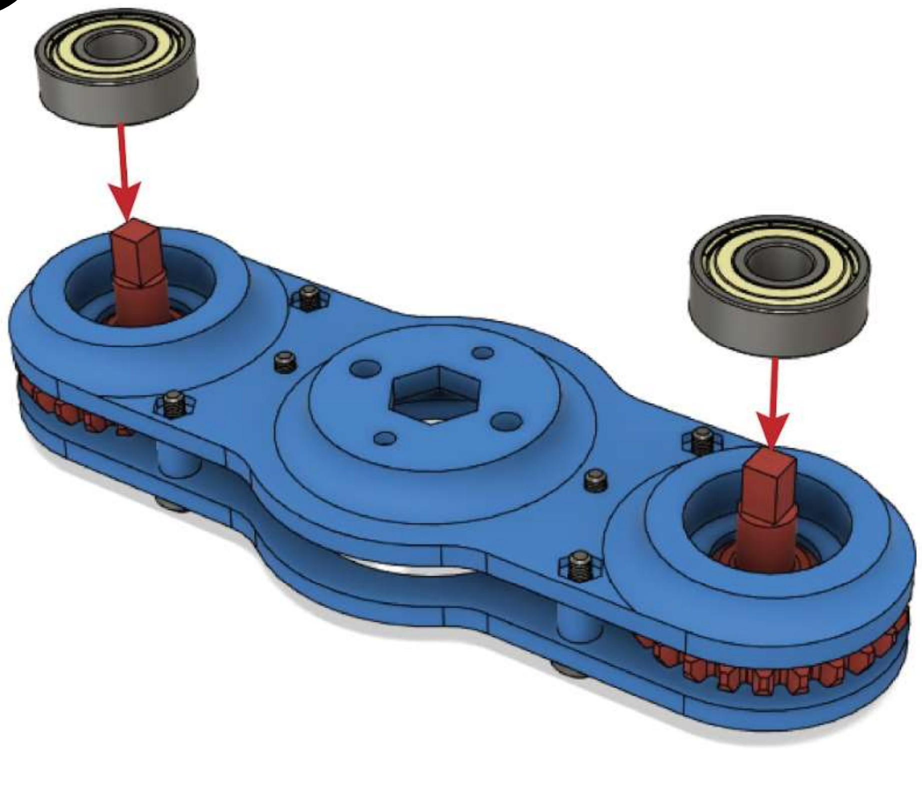

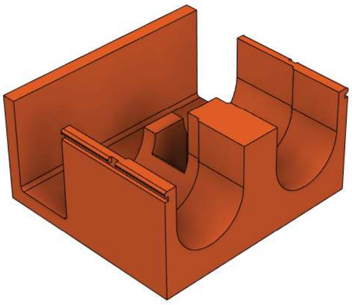

2 ×

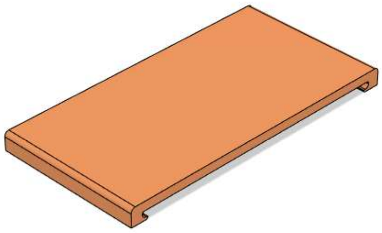

2 ×

**10**

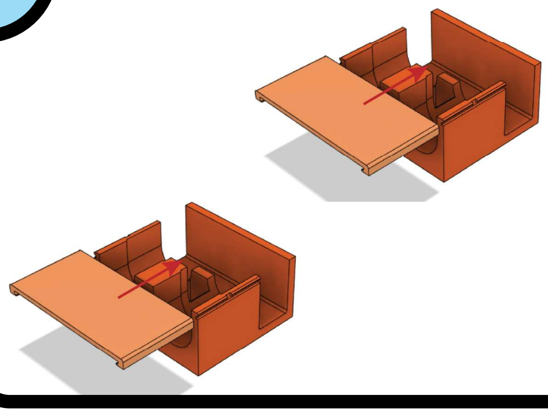

**11**

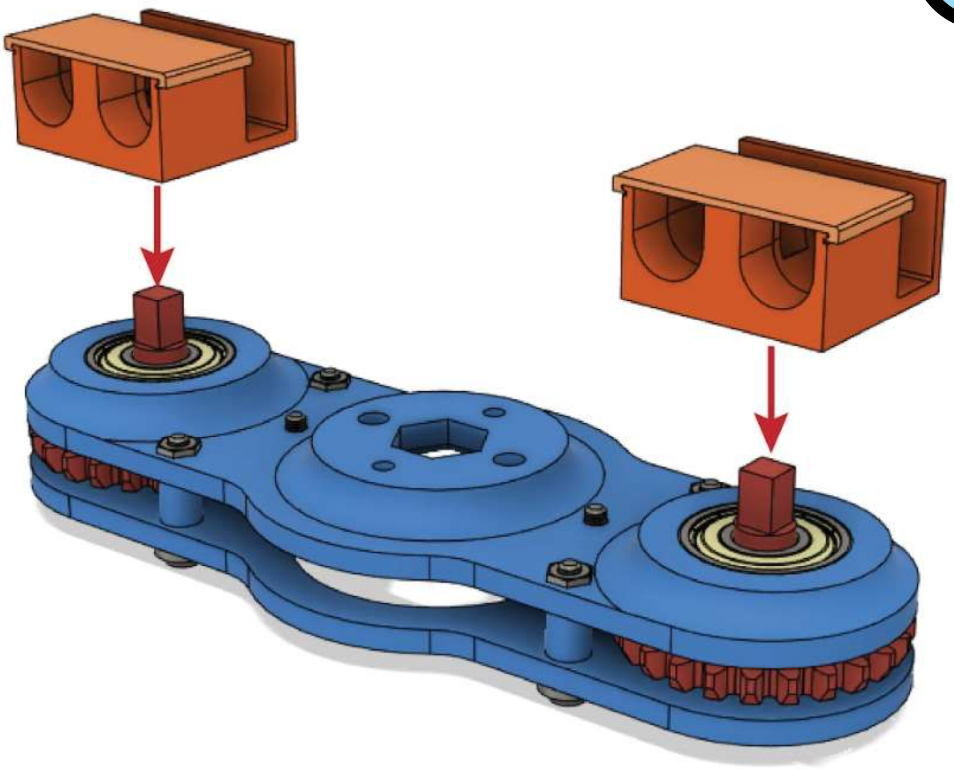

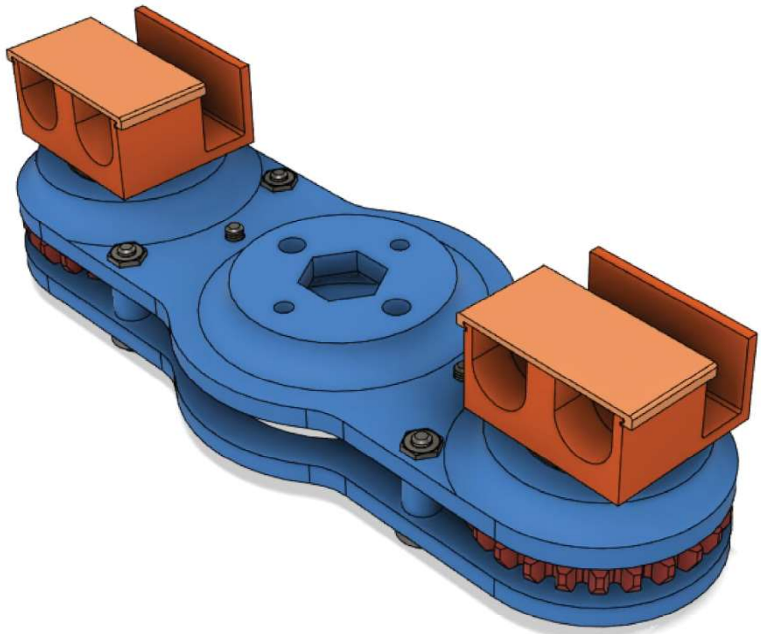

### Instructions for Assembly
### Centrifuge Attachment

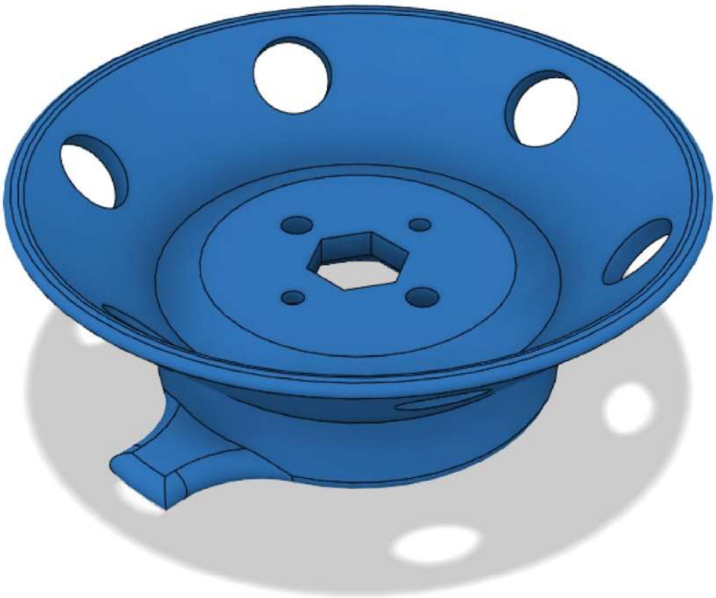

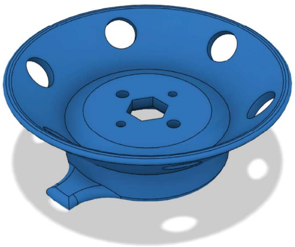

1 x

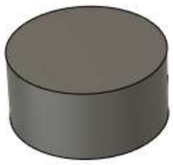

4 x

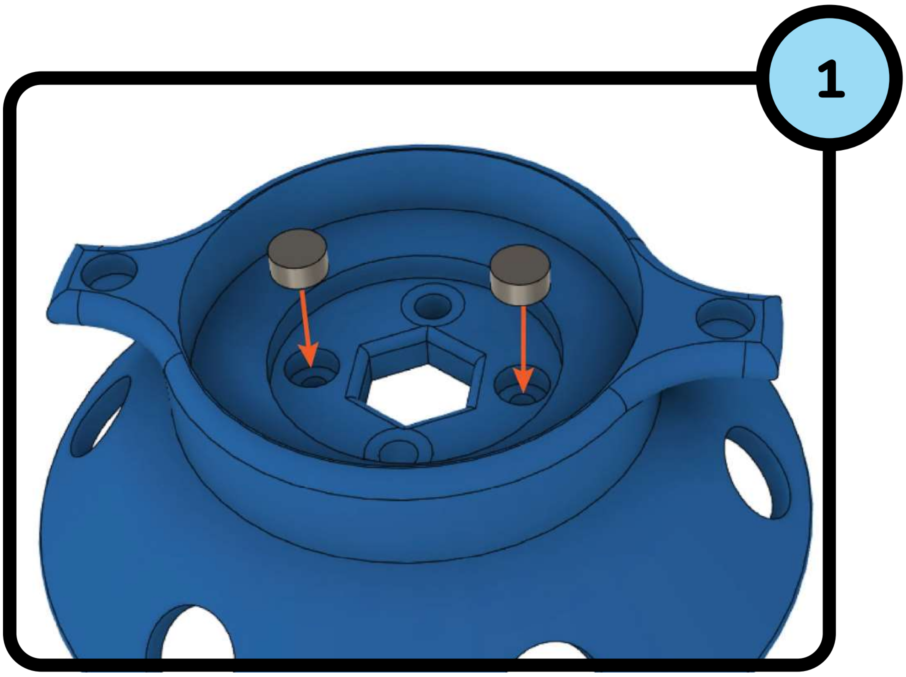


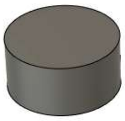

2 x

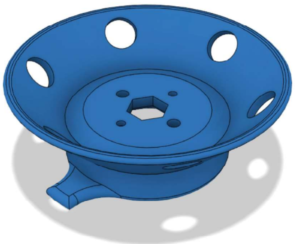

1 x

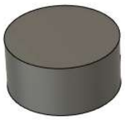

2 x

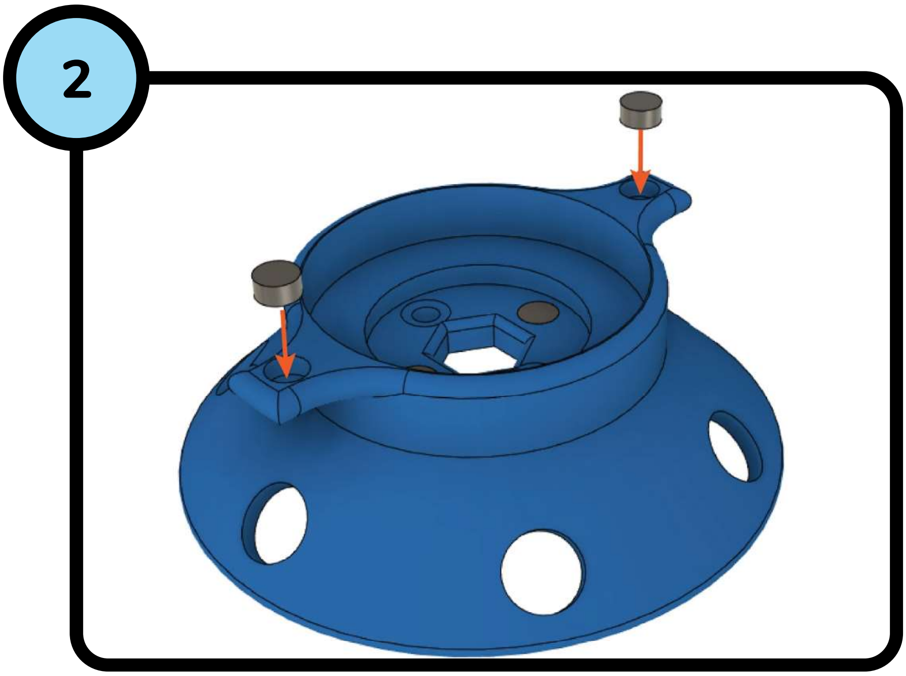


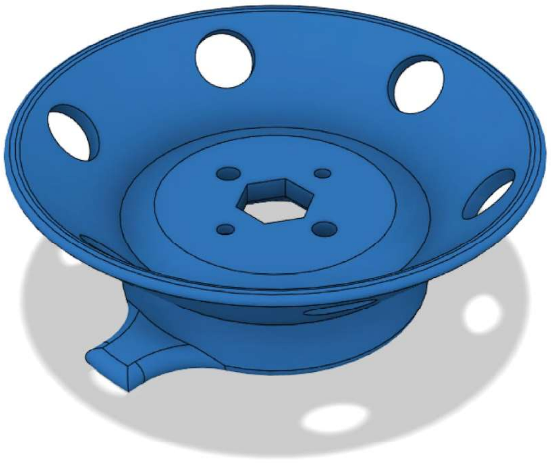

### Instructions for Assembly
### OpenCell Base

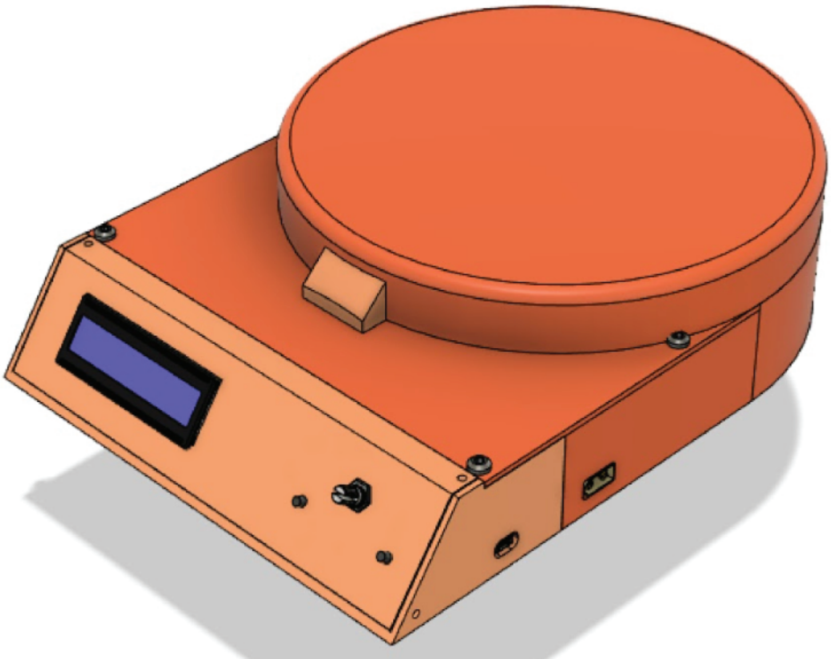

| | | | |
|---|---|---|---|
| 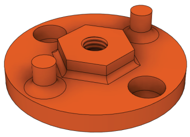 1 x | 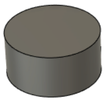 7 x | 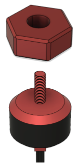 1 x | 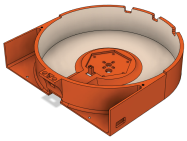 1 x |
| 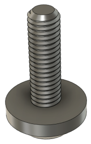 6 x m3x10mm | 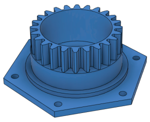 1 x | 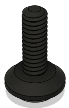 5 x motor screw | 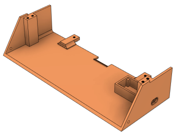 1 x |
| 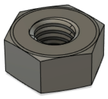 6 x | 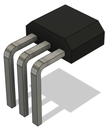 2 x | 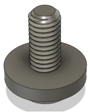 4 x m3x8mm | 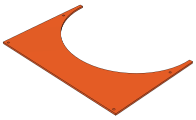 1 x |
| 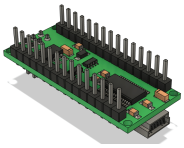 1 x | 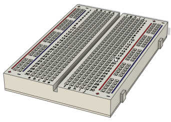 1 x | 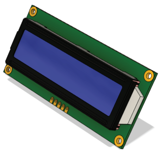 1 x | 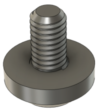 12 x m3x4mm |
| 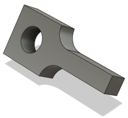 2 x | 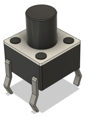 2 x | 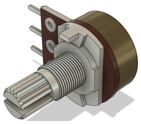 1 x | 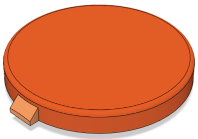 1 x |

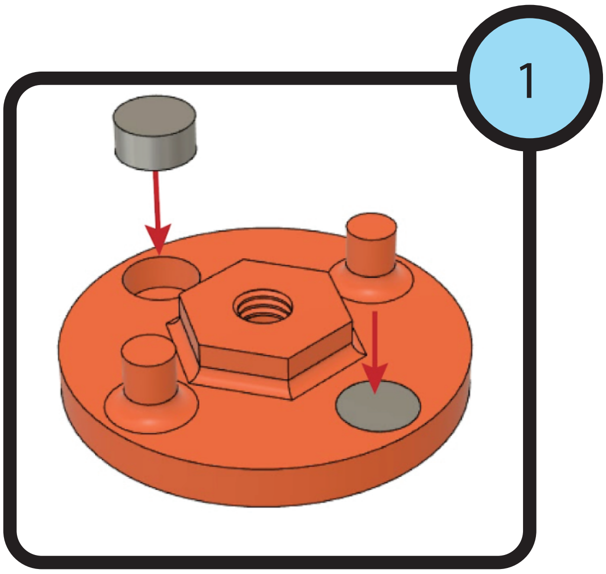


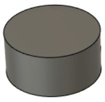

2 x

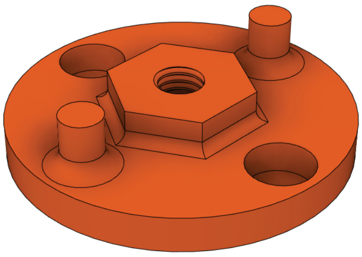

1 x

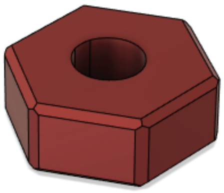

1 x

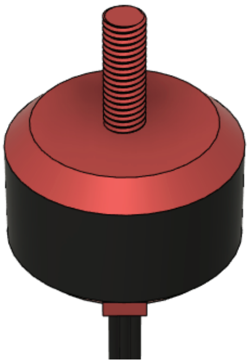

1 x

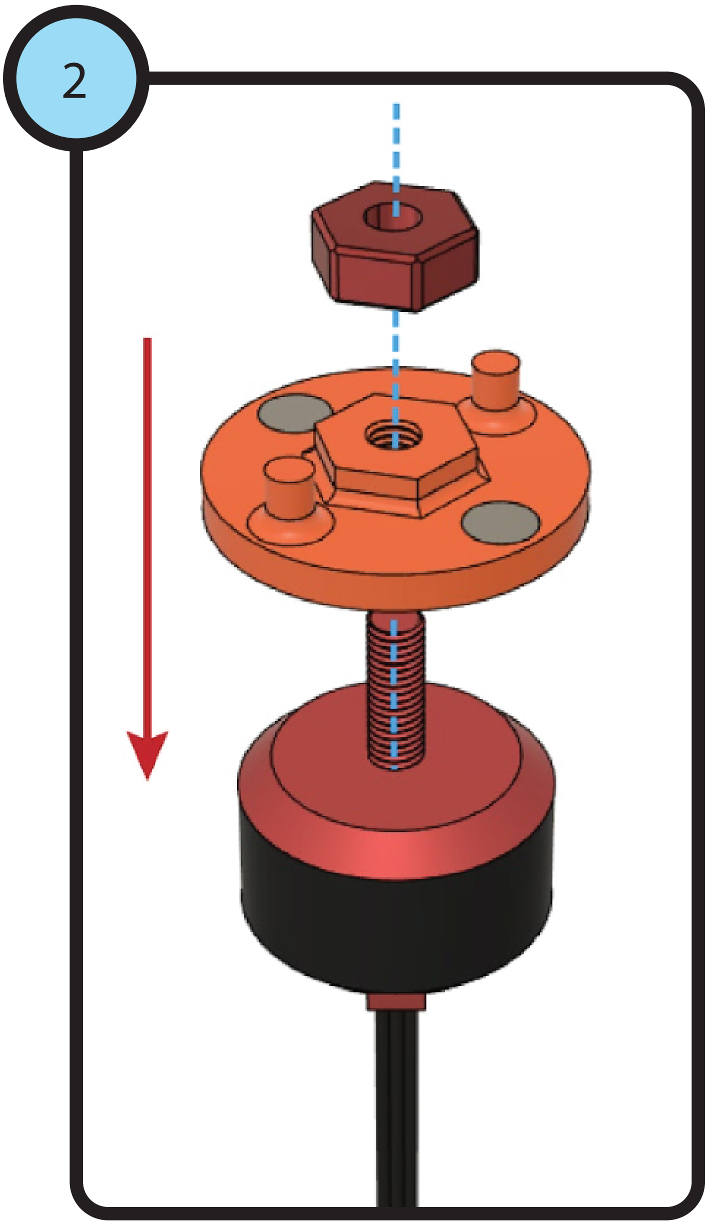

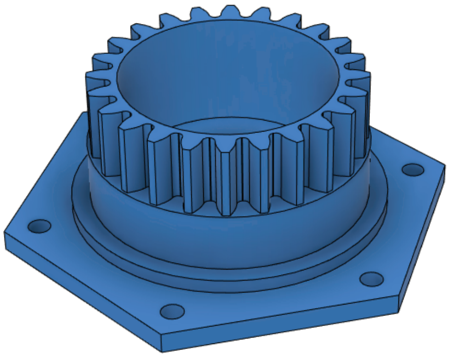

1 x

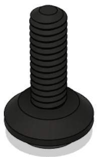

5 x
motor screw

3

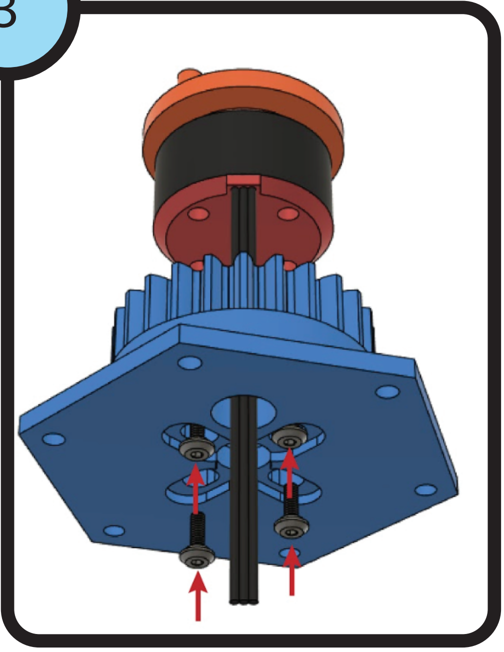

4

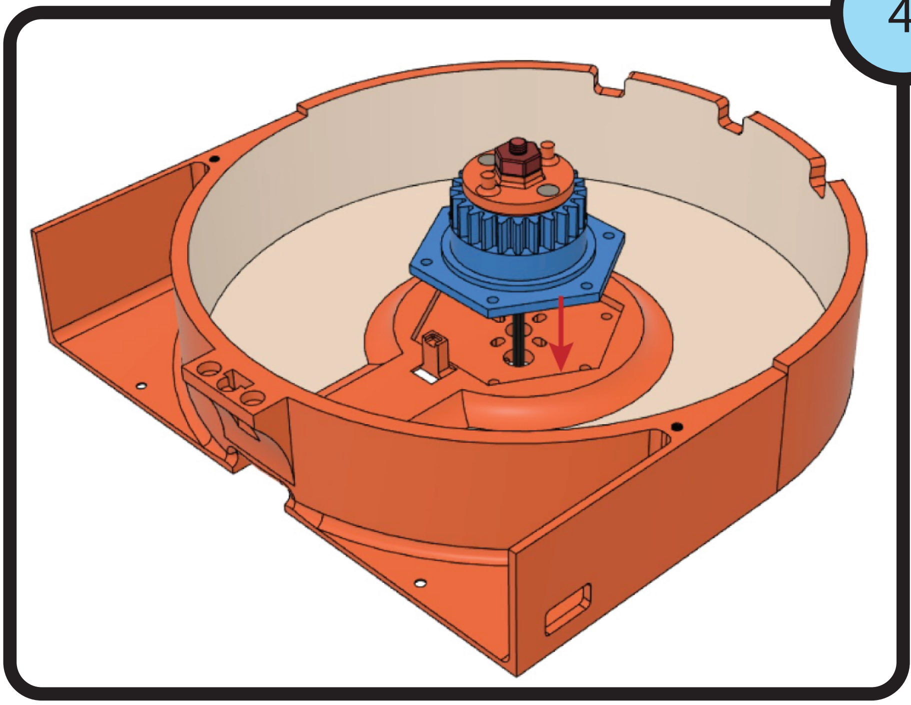

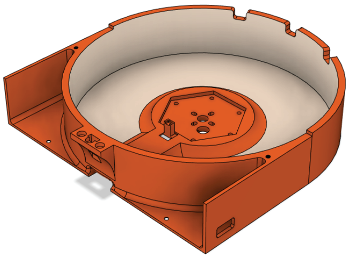

1 x

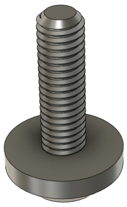

6 x
m3x10mm

5

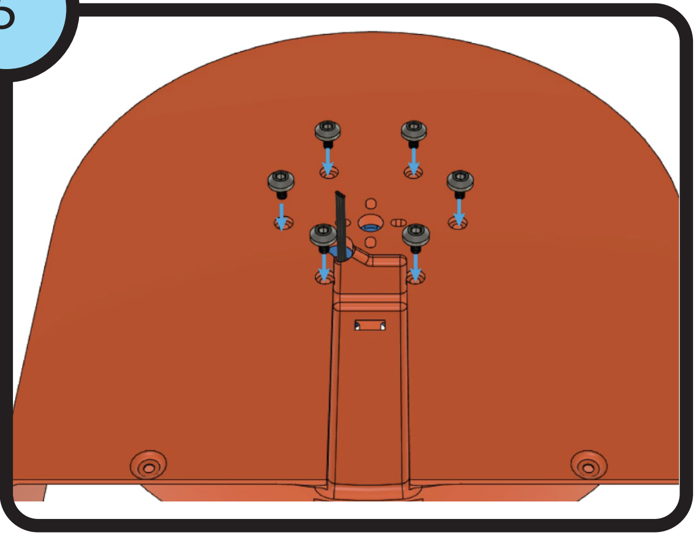

6

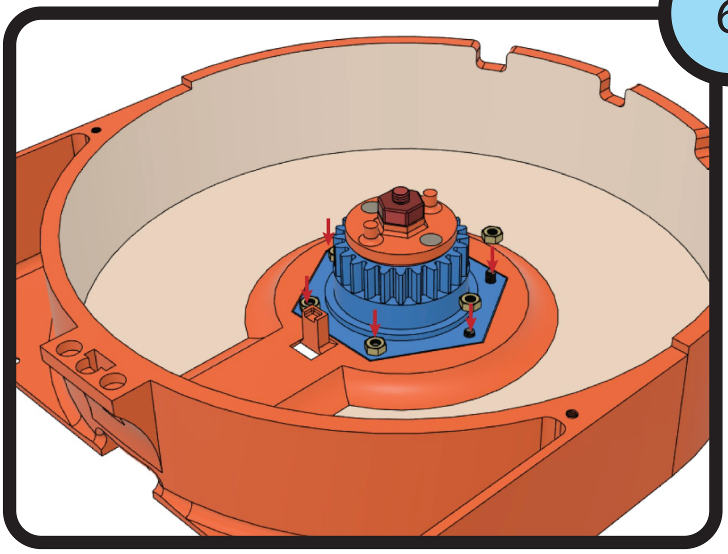

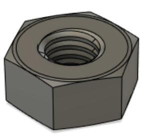

6 x

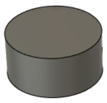

2 x

7

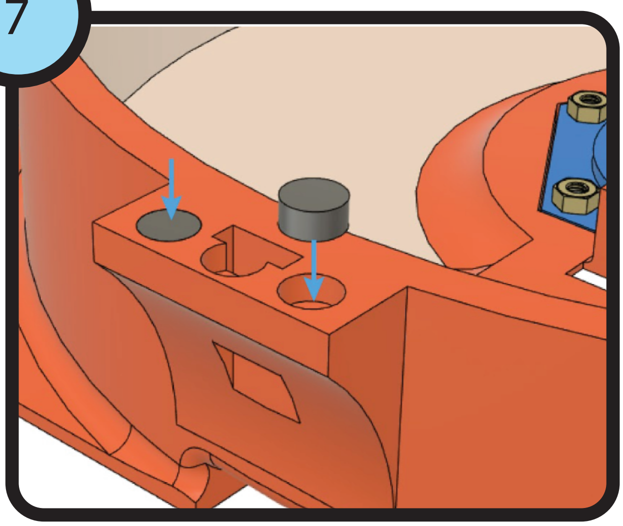

8

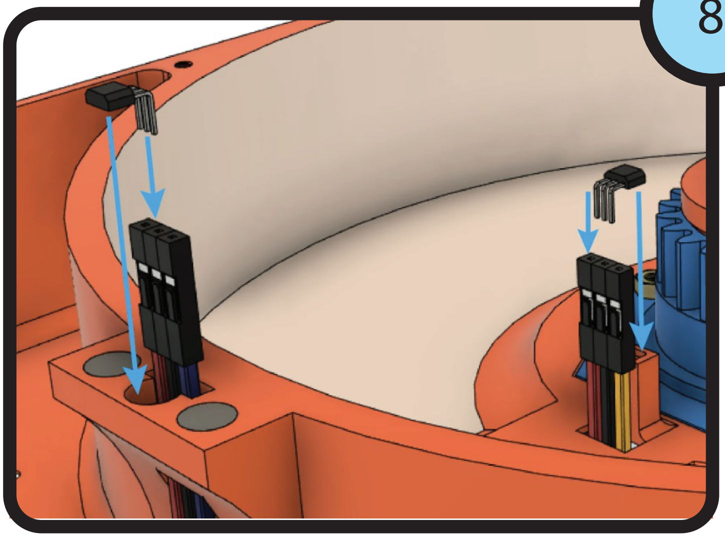

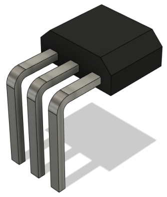

2 x

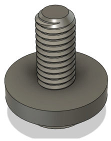

2 x
m3x8mm

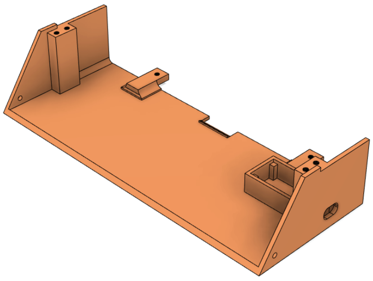

1 x

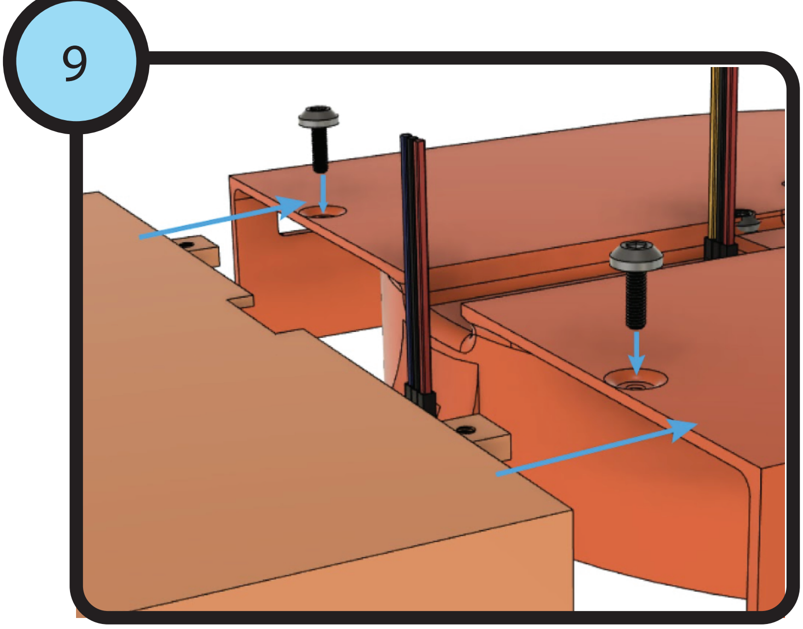


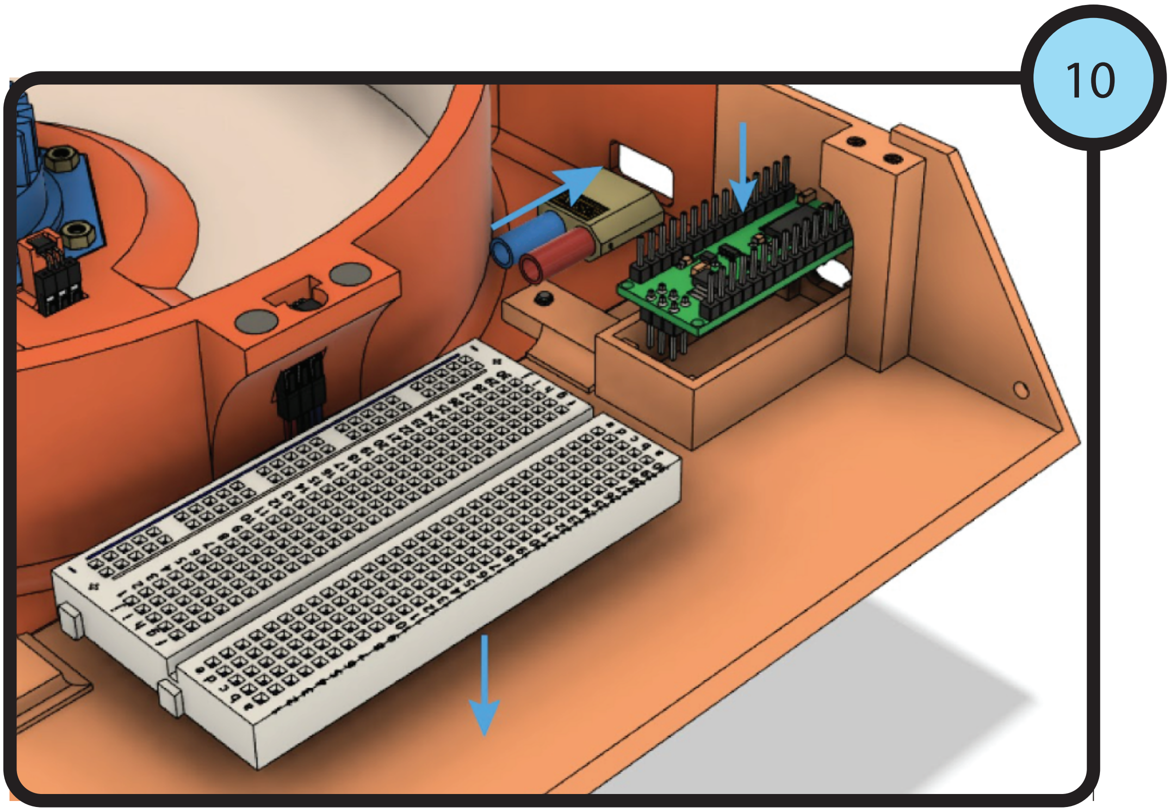


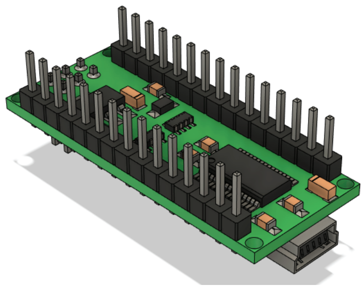

1 x

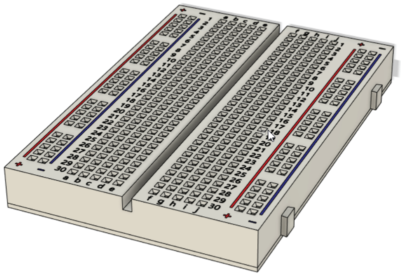

1 x

## 11

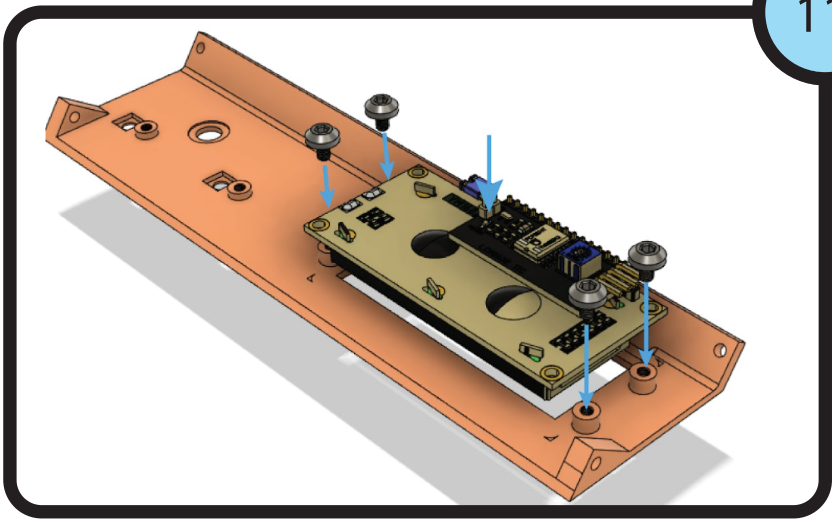

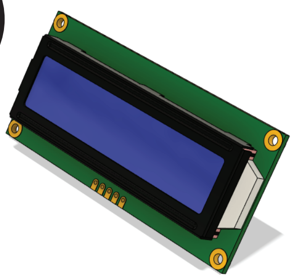

1 x

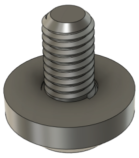

4 x
m3x4mm

## 12

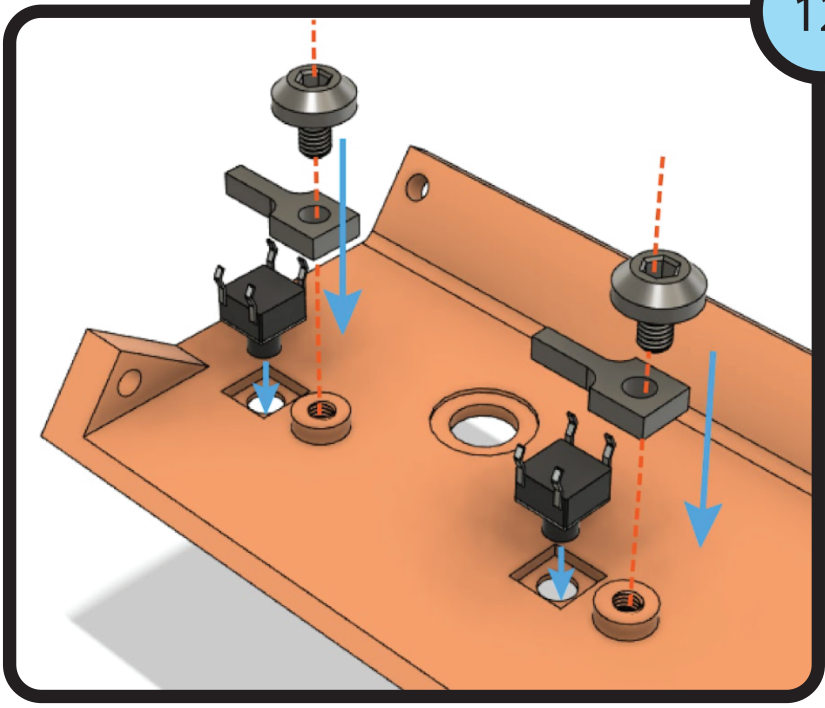

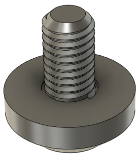

2 x
m3x4mm

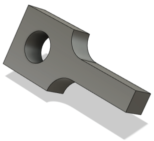

2 x

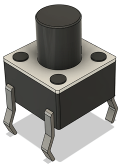

2 x

## 13

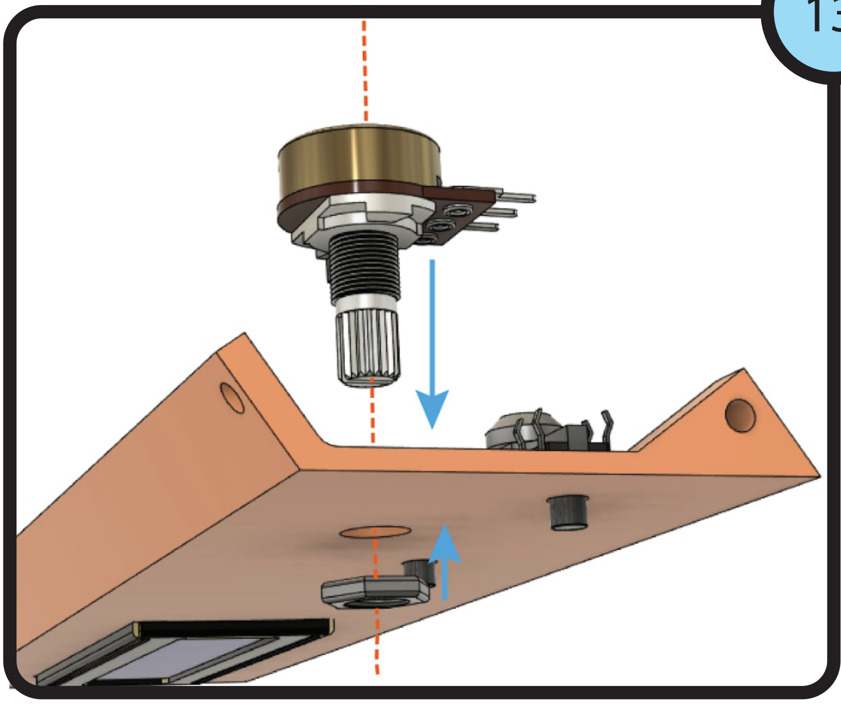

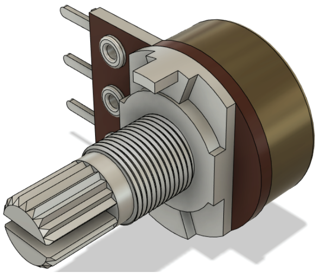

1 x

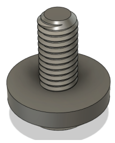

2 x
m3x8mm

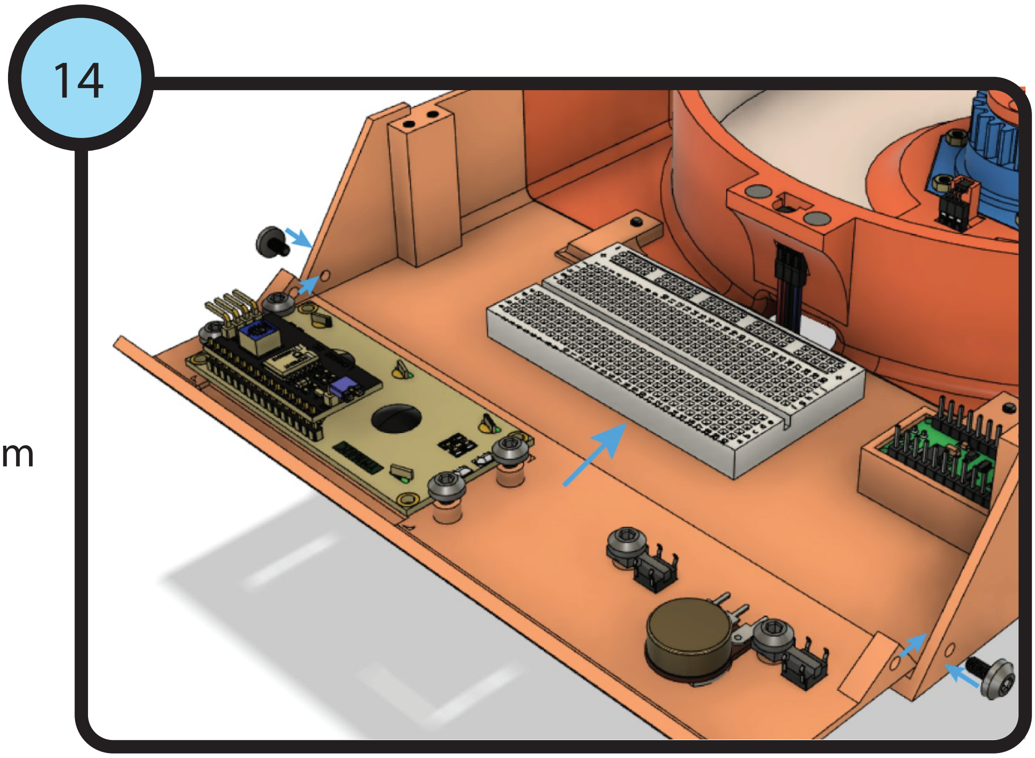


Note: all electrical connections can be found
in the wiring diagram (S1 Figure)

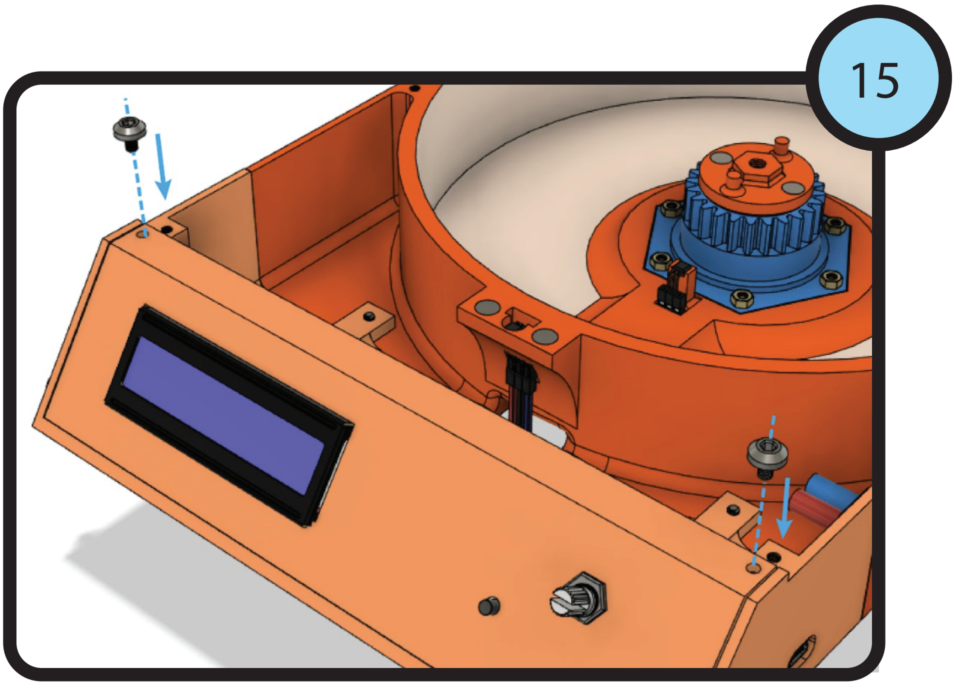


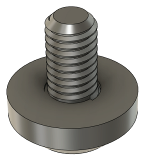

2 x
m3x4mm

## 16

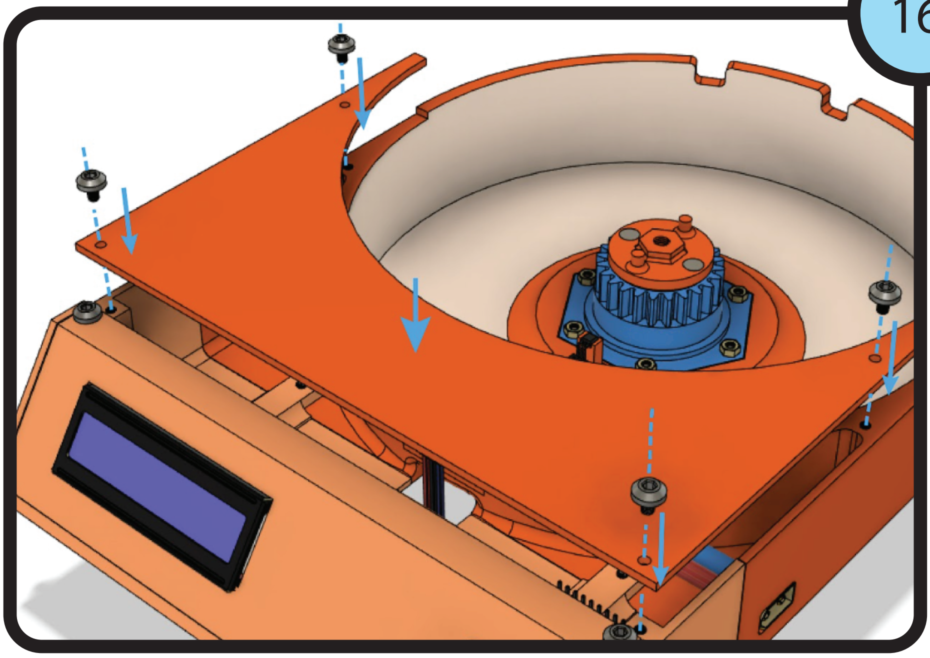

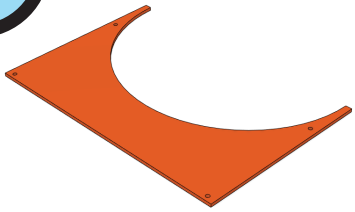

1 x

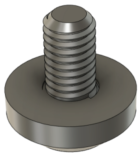

4 x
m3x4mm

## 17

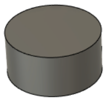

3 x

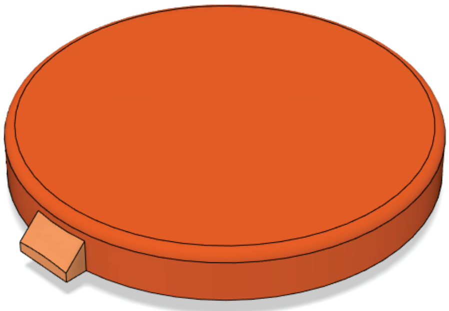

1 x

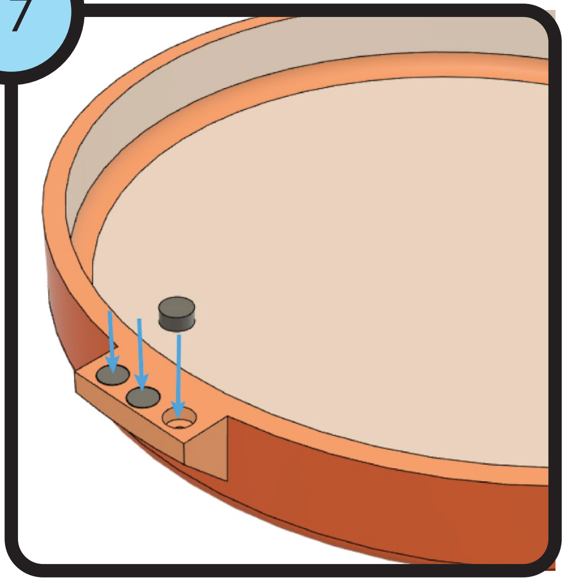

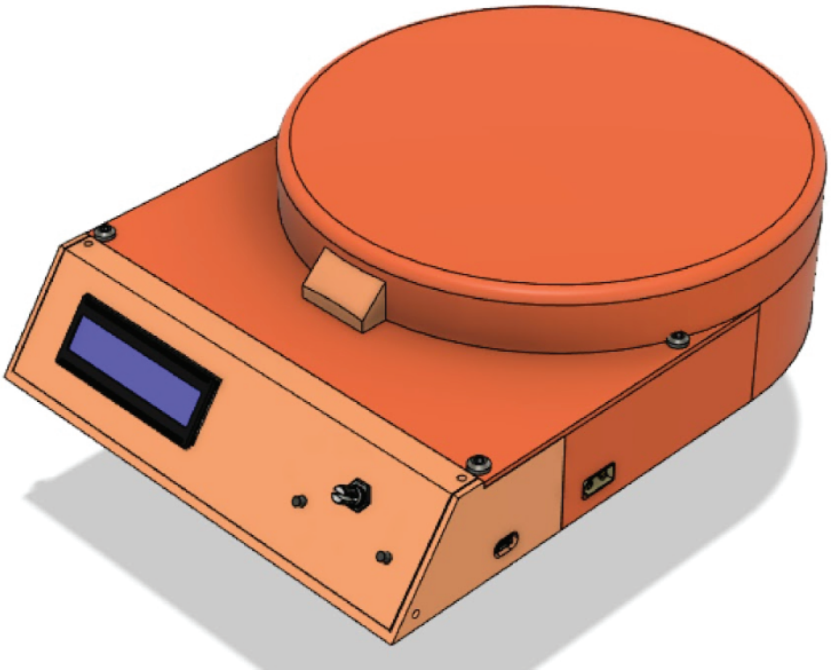
